# Classification and functional characterization of regulators of intracellular STING trafficking identified by genome-wide optical pooled screening

**DOI:** 10.1101/2024.04.07.588166

**Authors:** Matteo Gentili, Rebecca J. Carlson, Bingxu Liu, Quentin Hellier, Jocelyn Andrews, Yue Qin, Paul C. Blainey, Nir Hacohen

## Abstract

STING is an innate immune sensor that traffics across many cellular compartments to carry out its function of detecting cyclic di-nucleotides and triggering defense processes. Mutations in factors that regulate this process are often linked to STING-dependent human inflammatory disorders. To systematically identify factors involved in STING trafficking, we performed a genome-wide optical pooled screen and examined the impact of genetic perturbations on intracellular STING localization. Based on subcellular imaging of STING protein and trafficking markers in 45 million cells perturbed with sgRNAs, we defined 464 clusters of gene perturbations with similar cellular phenotypes. A higher-dimensional focused optical pooled screen on 262 perturbed genes which assayed 11 imaging channels identified 73 finer phenotypic clusters. In a cluster containing USE1, a protein that mediates Golgi to ER transport, we found a gene of unknown function, C19orf25. Consistent with the known role of USE1, loss of C19orf25 enhanced STING signaling. Other clusters contained subunits of the HOPS, GARP and RIC1-RGP1 complexes. We show that HOPS deficiency delayed STING degradation and consequently increased signaling. Similarly, GARP/RIC1-RGP1 loss increased STING signaling by delaying STING exit from the Golgi. Our findings demonstrate that genome-wide genotype-phenotype maps based on high-content cell imaging outperform other screening approaches, and provide a community resource for mining for factors that impact STING trafficking as well as other cellular processes observable in our dataset.

## Introduction

The cyclic-GMP-AMP (cGAMP) synthase (cGAS)/STimulator of Interferon Genes (STING) pathway plays a key role in anti-pathogen defense, anti-tumor responses, neurodegenerative disease, and autoimmunity^1^. Mechanistically, cGAS-mediated cGAMP production in response to pathogen-derived dsDNA drives innate immune responses that are beneficial to the host. In addition, recognition of mislocalized self-DNA of nuclear or mitochondrial origin drives immunity against tumors^2^. Dysregulation of this pathway leads to disease including development of autoinflammatory and neurodegenerative disease due to steady-state or ligand induced STING overactivation^3,4^.

Intracellular STING trafficking is essential for regulation of downstream responses. In the absence of ligand, STING is principally localized on the endoplasmic reticulum (ER) in the form of homodimers^5^. cGAMP ligation to STING leads to oligomerization of the sensor, which in turn activates STING trafficking from the ER to the Golgi. STING is palmitoylated at the Golgi leading to clustering and an increase in signaling^6,7^, and subsequently ubiquitination of the sensor^8^. STING then traffics to the trans-Golgi network (TGN), where it interacts with TANK-binding kinase 1 (TBK1) to phosphorylate IRF3 and produce type I interferon (IFN)^7^. Signaling shutdown is achieved by ESCRT-mediated degradation of ubiquitinated STING at the lysosome^8–10^. STING oligomerization has also been shown to drive autophagy activation, an ancestral activity of the pathway important in pathogen clearance^11^. Activation of STING-dependent autophagy relies on a channel formed in the transmembrane domain of STING dimers, leading to pH imbalance in the Golgi compartment^12^. Finally, STING has also been shown to activate the NF-κB pathway to induce pro-inflammatory cytokines, even though the mechanisms leading to this response are less clear^13^.

In addition to cyclic di-nucleotide mediated trafficking, we and others have shown that cGAS basal activity primes STING, leading to a homeostatic degradative flux, and post-Golgi interruption of this flux leads to spontaneous activation of the sensor^8,14^. This is a process of particular physiological relevance, since mutations in genes regulating STING trafficking have been found in patients with autoinflammatory diseases. Mutations in the COP-I subunit COPA found in patients with COPA syndrome lead to STING accumulation at the Golgi, resulting in autoinflammation in patients^15–19^. Similarly, mutations in two genes driving neurodegenerative diseases, C9orf72 for Amyotrophic Lateral Syndrome (ALS) and NPC1 for Niemann-Pick Disease, impair STING degradation at the lysosome, inducing heightened or constitutive STING-dependent responses and resulting in disease^20,21^. Maintaining homeostasis of the STING pathway is therefore central to avoiding exacerbated activation of inflammatory responses. On the other hand, blocking STING degradation has been shown to be beneficial in the context of antitumor immunity, since STING-dependent type I IFN, central to activation of the immune system against tumors, can be boosted^14,22^. Understanding regulators of STING trafficking is therefore important both in the context of autoinflammation and antitumor immunity.

In order to identify genes regulating STING trafficking, we previously performed flow-based CRISPR screens and proximity-ligation proteomics that led us to characterize ESCRT as a regulator of STING degradation at the endosome^8^. The screen used a terminal readout of STING degradation at the lysosome, but STING trafficking is a dynamic and multifaceted process that can be followed intracellularly via microscopy. Here, we set out to associate genetic perturbations with specific intracellular trafficking phenotypes upon STING activation. To achieve this goal at the genome-wide level, we took advantage of Optical Pooled Screening (OPS)^23^, a method that enables high-throughput image-based pooled genetic screens in tens of millions of cells^24,25^. We show that OPS outperforms previous STING screens relying on one-dimensional flow cytometric or fitness read-outs using a meta-analysis approach^26^. In addition, this high-content screen allows for further interpretation directly from screening data. Indeed, using dimensionality reduction and unsupervised clustering^24,25,27^, we identify groups of genes that similarly affect STING trafficking when knocked out and we show that this dataset is useful for assigning functional roles to previously uncharacterized genes. We focused on two groups of genes regulating STING degradation at distinct stages and found that the HOPS complex is required for signaling shutdown at the endolysosome. Similarly, we show that the GARP complex subunit VPS52 and the RAB6 GEF RIC1/RGP1 complex drive STING degradation by promoting its exit from the Golgi. Overall, by performing high-content OPS, we provide a resource to the community to guide identification of genes regulating STING biology.

## Results

### A genome-wide optical pooled screen efficiently retrieves genes involved in STING trafficking

In order to identify genetic regulators of STING subcellular trafficking, we used optical pooled screening (OPS) in HeLa-TetR-Cas9 cells (**Fig. 1a**), and induced Cas9 activity for 7 days through doxycycline administration^28^. HeLa cells were transduced with STING-mNeonGreen (mNG) to monitor STING localization and flow sorted twice to select for cells with similar reporter expression levels. We used a library of sgRNAs targeting ∼20,000 genes with 4 sgRNAs/gene, along with 454 non-targeting sgRNAs (ntgRNAs) previously described^24^. Following library transduction and doxycycline administration, cells were stimulated with cGAMP for 4 hours. Subcellular compartments involved in STING trafficking (Golgi - GM130, and endolysosomes - CD63) were imaged in addition to p62, an autophagy receptor that accumulates in response to STING activation and has been shown to interact with STING^29^ (**Fig. 1b**). As expected, cGAMP induced STING translocation from the ER to the Golgi apparatus and endolysosomal compartments (**Fig. 1b**). In addition to the 4 hour stimulation time-point used for the perturbation screen, two unperturbed control conditions (cells that received a non-targeting sgRNA) were acquired as comparators: 1) unstimulated cells as a baseline for STING protein distribution at steady-state and 2) cells at a later time-point, 5 hours after cGAMP stimulation, to assess further changes in the distribution and degradation of STING.

**Figure 1.**
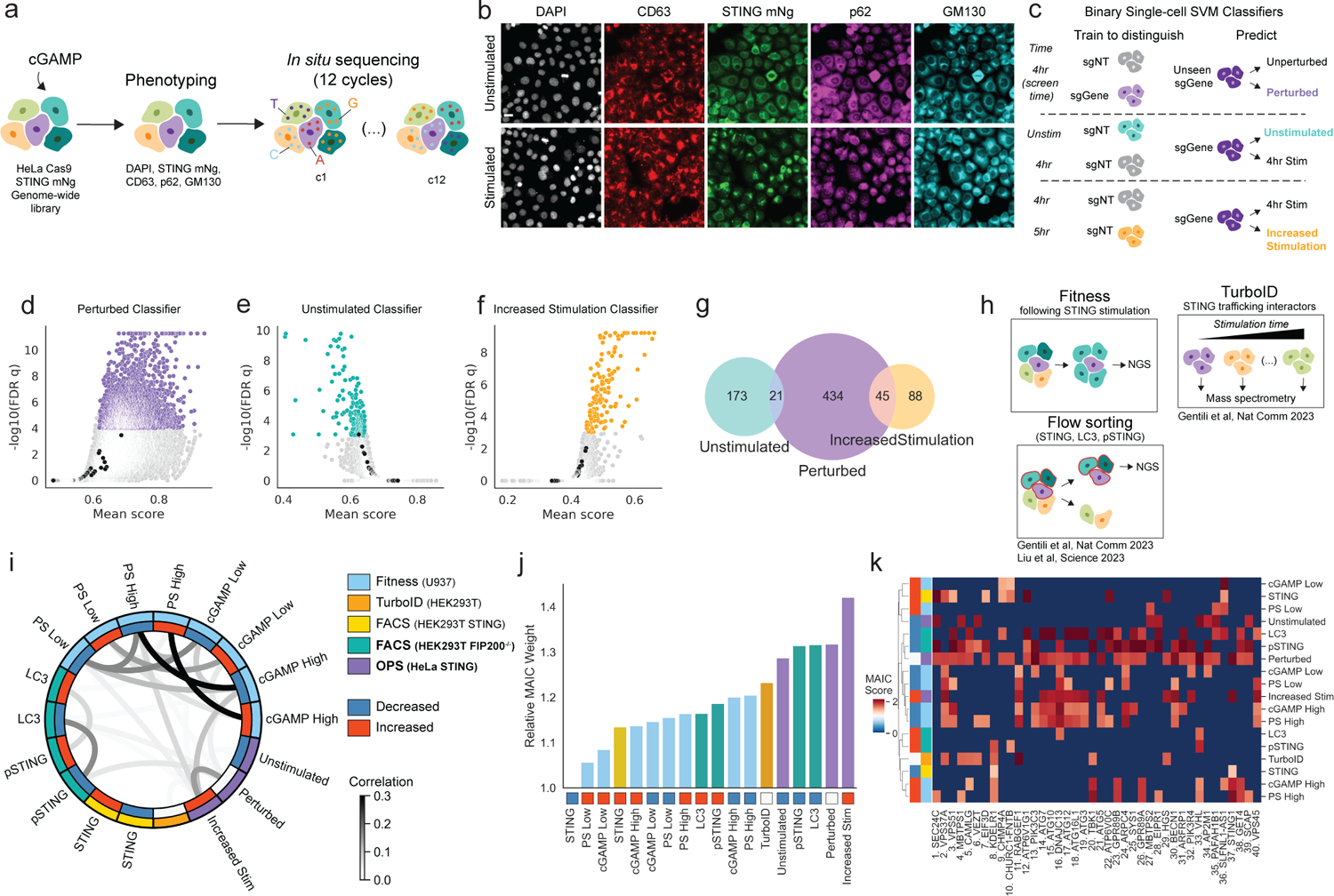
A genome-wide optical pooled screen identifies genes regulating STING trafficking. **(A)** Genome-wide optical pooled screening workflow for high-throughput STING screens. **(B)** Example fields of view of HeLa cells from the optical pooled screen either unstimulated (top) or stimulated (bottom) with cGAMP for 4 hours. Scale bar 20 µm. **(C)** Schematic of single-cell support vector machine (SVM) classifiers used to call hits from the genome-wide optical pooled screen using features from the p62 and STING channels. 4hrs indicates cGAMP-stimulated cells that received the genome-wide KO library. 5hrs refers to control cells with non-targeting sgRNAs stimulated with cGAMP for 5 hours. **(D-F)** Volcano plot of hits from the perturbed vs unperturbed classifier **(D)**, unstimulated classifier **(E)**, increased stimulation classifier **(F)**. Black dots indicate individual non-targeting sgRNAs. **(G)** Venn diagram showing overlap of top 500 significant hits from each classifier. **(H)** Genome-wide fitness and flow cytometry screening outlines and mass spectrometry STING interaction screen. **(I)** Chord diagram showing Pearson correlation between MAIC scores for the top 500 significant hits from each of the STING datasets. Same experimental batch performed on the same day highlighted by the same color in the outer ring. **(J)** Weighting score for each of the STING datasets provided by the MAIC algorithm. The weighting score given to a dataset is proportional to the average score of the genes belonging to this dataset and genes appearing in multiple datasets score more highly. **(K)** Scores for each of the top 40 genes in each screening condition.

Following imaging and *in situ* sequencing of 45 million cells with 80,000 perturbations (**Fig. S1a**), we defined the phenotype of each cell based on thousands of features extracted from the images, including intensity, marker co-localization, subcellular variation, texture, and shape (using CellProfiler^30^, scikit-image^31^, and mahotas^32^) (**Table S1**). Using all features derived from the STING or p62 channels (**Fig. S1b**) and including information from other channels such as colocalization with CD63 or GM130, we developed three independent SVM classifiers (**Table S2**) to determine which 4-hour stimulated cells in the screen display phenotypes that are: 1) significantly different from cells with non-targeting guides (but still cGAMP-stimulated) and thus defining 500 gene hits based on the mean of 4 guides (‘perturbed’ classifier; **Fig. 1d**); 2) similar to those of control unstimulated cells, identifying 194 genes required for STING exit from the ER/Golgi (‘unstimulated’ classifier; **Fig. 1e, S1c**); 3) similar to control cells that are stimulated for 5 hours with cGAMP, finding 133 genes that may accelerate STING trafficking or lead to STING accumulation in the endolysosomal compartment (‘stimulated’ classifier; **Fig. 1f, S1c**). The two latter classifiers identified genes not found by the first classifier (**Fig. 1g**), likely because of increased power for detecting specific stimulation conditions thanks to classifier training and feature selection on these conditions. The classifiers reported known genes: for instance, SEC24C, which is critical for STING exit from the ER^11^ and should therefore result in cells appearing unstimulated when knocked out, was one of the top 5 hits in the unstimulated classifier and also a hit in the perturbed classifier, but not in the increased stimulation classifier. On the other hand, VPS37A, which results in increased STING accumulation when knocked out^8^, was one of the top 15 hits in the increased stimulation classifier and scored in the perturbed classifier but not in the unstimulated classifier. In total, the optical screen identified 695 gene hits at FDR q <= 0.001 in at least one model.

We next combined the results of the optical screen with other non-imaging-based genome-wide genetic screens that we carried out to find regulators of STING activation based on: 1) STING-dependent cell fitness in U937 cells (reported here); 2) STING levels in HEK293T^8^; 3) LC3B levels^12^ and 4) pSTING levels (reported here), with the latter two screen in HEK293T FIP200 KO cells to reduce background levels of canonical autophagy^12^ (**Fig. 1h-S1e**). To complement these genetic screens, we also included 132 genes encoding proteins that we found in physical proximity to STING at 4 timepoints after activation using proximity-ligation proteomics^8^.

We then examined the correlation of hit genes across the optical screen and these genetic and proteomic studies (**Fig. 1i**). In addition to higher Pearson correlations of relative meta-analysis by information content (MAIC)^26^ scores for screens performed at the same time, we observed that genes for which knockout increased cell death were correlated with genes found in the increased stimulation classifier, while those that decreased cell death, pSTING, or STING intensity levels by flow cytometry were often included in our perturbed classifier. Using our previously described MAIC approach to compare the power of each screen to find true positives (across 18 gene lists and 9 screens), the top two performers were optical pooled screening classifiers (increased stimulation and perturbed) and the remaining OPS classifier - unstimulated - was ranked 5th (**Fig. 1j**). Therefore, despite the high quality of the previous datasets that revealed several novel STING regulators and functions^8,12^, our high-content image-based OPS dataset resulted in more informative gene lists than previous approaches. By integrating multiple screens performed in distinct biological models, we developed a consensus list of 1225 genes that scored in at least two distinct screens (**Table S3**), yielding increased confidence in the broad biological relevance of these genes to the STING pathway. Indeed, even among the top 40 genes, many of which are well-established regulators of the STING pathway, no gene scored in all screening settings (**Fig. 1k, S1d**), revealing the importance of dataset integration for prioritization of high-confidence genes.

### Unsupervised clustering and dimensionality reduction reveal genes with similar effects on STING trafficking

We further explored the biological functions of these 1225 consensus genes by performing unsupervised analyses on the image-based features obtained from our high-content genome-wide optical pooled screen. To do this, we performed dimensionality reduction on thousands of gene-level mean features using the PHATE algorithm^33^ and performed Leiden clustering^34^ to identify 464 clusters of genes of interest (**Fig. S2a-b, Table S4**). To further infer the functions of genes in these clusters, we identified clusters that were enriched for hits derived using our unstimulated classifier (Fig. 2a). Among the clusters that scored as significant, cluster 1 included a number of ER membrane genes, including two known regulators of STING ER exit, SEC24C^11^ and SCAP^35^, as well as other ER-localized genes not yet shown to regulate STING such as MBTPS1 and MBTPS2 that were recently reported to act with SCAP to regulate NF-kB^36^. We also observed a cluster of dynein/dynactin factors (cluster 18); these complexes have previously been associated with ER exit but not shown to be required for STING activation^37^. Finally, we observed that the GARP complex delayed STING activation at a later stage, causing cells to look significantly more like unperturbed cells at 4 hours than unperturbed cells at a longer 5 hour stimulation timepoint.

**Figure 2.**
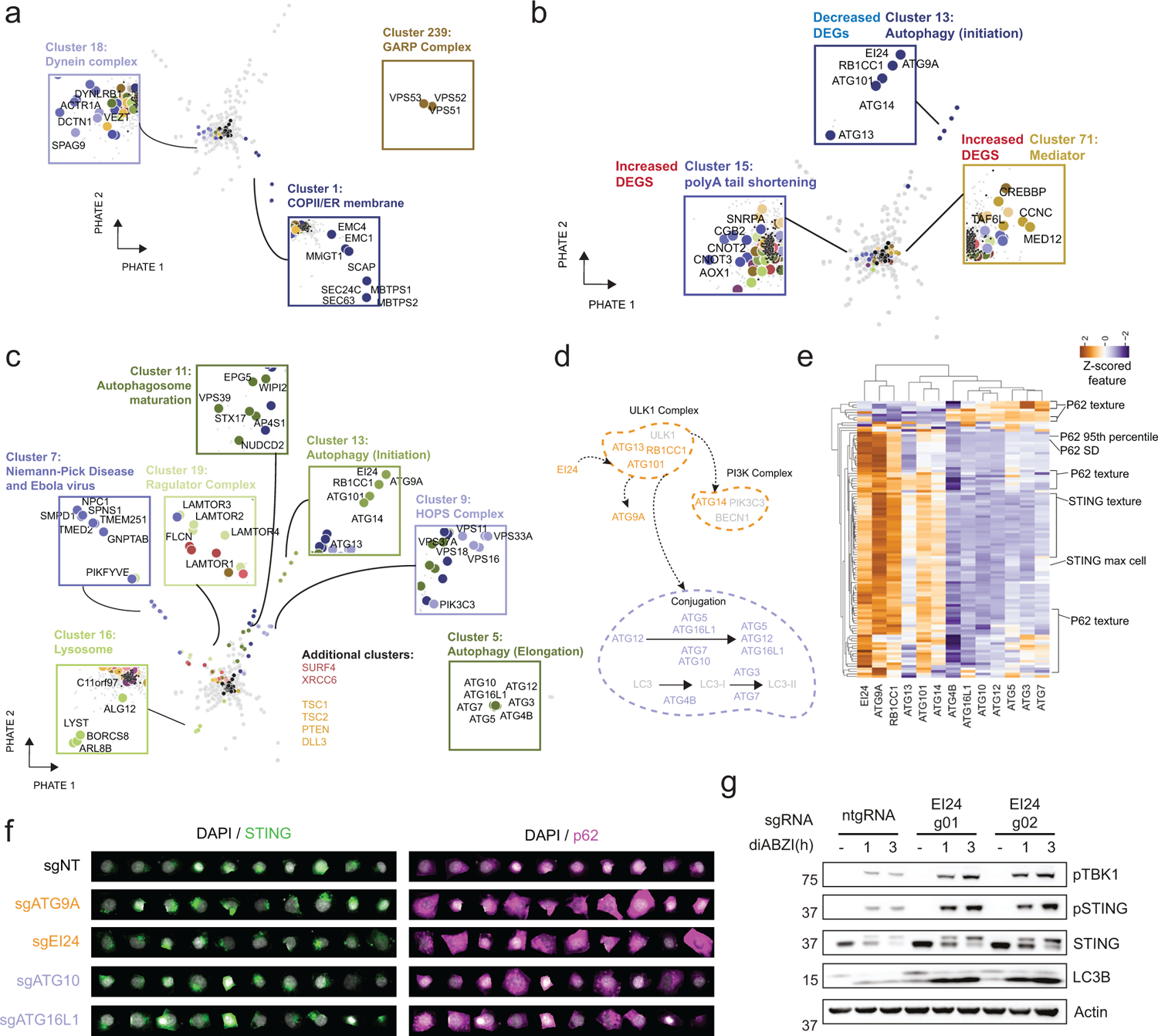
Unsupervised clustering identifies groups of genes that similarly affect STING trafficking. **(A)** PHATE plot highlighting top clusters significantly enriched for genes that are less stimulated because they are either more similar to unstimulated non-targeting control cells (clusters 1, 18 - unstimulated classifier) or significantly more similar to non-targeting control cells at 4 hours than at 5 hours (cluster 239). **(B)** PHATE plot highlighting clusters significantly enriched for genes that have a high or low number of differentially expressed genes in the genome-scale Perturb-seq dataset. **(C)** PHATE plot highlighting top clusters significantly enriched for genes that are more stimulated than non-targeting control cells (increased stimulation classifier). **(D)** Autophagy diagram highlighting genes identified in cluster 13 (orange) or 5 (purple). **(E)** Hierarchically clustered heatmap (clustered using seaborn clustermap with method=’average’, metric=’euclidean’), showing features differentiating genes from the two autophagy clusters in (D). **(F)** Single-cell images of STING and p62 channels randomly selected from non-targeting controls and KO of selected autophagy factors from the genome-wide OPS. **(G)** Immunoblot of the indicated proteins in control (ntgRNA) or EI24 KO BJ1 fibroblasts stimulated with 1µM diABZI for the indicated times. One representative blot of n=3 independent experiments.

To test if the hits identified in our screens impacted transcription in resting cells, potentially pointing to perturbation effects acting indirectly on STING, we integrated our dataset with the genome-scale Perturb-seq dataset^38^. By doing so, we identified very few clusters with enrichment (clusters 15, 71) for perturbations that led to hits in the genome-scale Perturb-seq screen with significant differentially expressed genes (Fig. 2b**, S2c**), increasing confidence that many of our hits do not affect general cellular processes and likely have a direct impact on STING trafficking. To further increase confidence in our clustering approach, we tested whether clusters at a significant distance from the ntgRNAs were enriched for protein-protein interactions (PPI) as defined in the CORUM dataset^39^ and confirmed that mean distance from ntgRNAs positively correlated with an enrichment in PPIs (**Fig. S2d**). Additionally, clusters containing more than one gene and less than 20% ntgRNAs showed significant enrichment for GO, Reactome, KEGG, CORUM and Jensen terms (**Fig. S2e**). This confirmed that our clustering approach identified biologically related groups of genetic perturbations.

Next, we identified clusters of genes that were enriched for genes with increased stimulation in our classifier (Fig. 2c). These include a cluster containing SURF4, a protein that when knocked out results in increased accumulation of STING at the Golgi^17^, the HOPS complex, and several clusters related to autophagy (5, 11, and 13). In fact, when examining two distinct stages of autophagy, initiation and conjugation, we found that our screen identified the majority of genes critical for these stages and that they clustered separately as cluster 5 (orange) and cluster 13 (purple) (Fig. 2d**-e**) mostly based on intensity and texture features from both the STING and p62 channels. Single-cell images indeed revealed distinct patterns for p62 and STING for autophagy initiation and conjugation factors (Fig. 2f). Notably, EI24, a gene involved in autophagy that was only recently revealed to be involved in initiation, clustered with other canonical initiation factors^40^. We therefore experimentally confirmed that EI24 knockout significantly affects STING signaling by knocking out this gene in hTERT-immortalized primary BJ1 fibroblasts. Via Western blotting, we observed an increase in LC3B lipidation, a reduction in STING degradation and a consequent increase in pSTING and pTBK1 downstream signaling, suggesting that, like ATG9A, which clustered with EI24 (Fig. 2c**-d**) and was shown to regulate STING trafficking^41^, EI24 is involved in STING signaling shutdown (Fig. 2g**, S2f**).

### Secondary screens further characterize genes involved in STING trafficking and signaling

To further confirm the effect of 262 top genes involved in STING trafficking, we performed follow-up screening in HeLa cells (**Table S5**) and BJ1 fibroblasts (**Table S6**), thus diversifying the cellular models studied (Fig. 3a**-c**). We imaged at higher resolution in (1) time, with unstimulated, 4 hour, and 6 hour post cGAMP treatment for HeLa, and 4 hours for BJ1 and (2) phenotype, by expanding to 11 imaging channels using the iterative immunophenotyping method 4i in HeLa^42^ (Fig. 3a**-c**). We visualized proteins relevant to STING signaling (STING, NF-κB p65, p62), STING trafficking compartments (Calnexin in the ER, GM130 in Golgi, EEA1 in endosomes), STING degradation (HGS, ATP6V1D), and cell state (DAPI/nuclei, actin, γH2AX) (example images for all channels in **Fig. S3a**). As with the genome-wide screen, we extracted thousands of features from each channel without background subtraction as well as with the addition of a rolling ball background subtraction step (Fig. 3c**, Fig. S3a**)^43^. In HeLa cells, STING colocalization with Calnexin decreased after stimulation with cGAMP, while STING colocalization with GM130, HGS and EEA1 increased (**Fig. S3b**). Additionally, the NF-κB subunit p65 showed increased signal intensity in DAPI positive nuclei upon cGAMP stimulation (**Fig. S3b**). These results recapitulated the expected responses and served to technically validate the 4i stainings.

**Figure 3.**
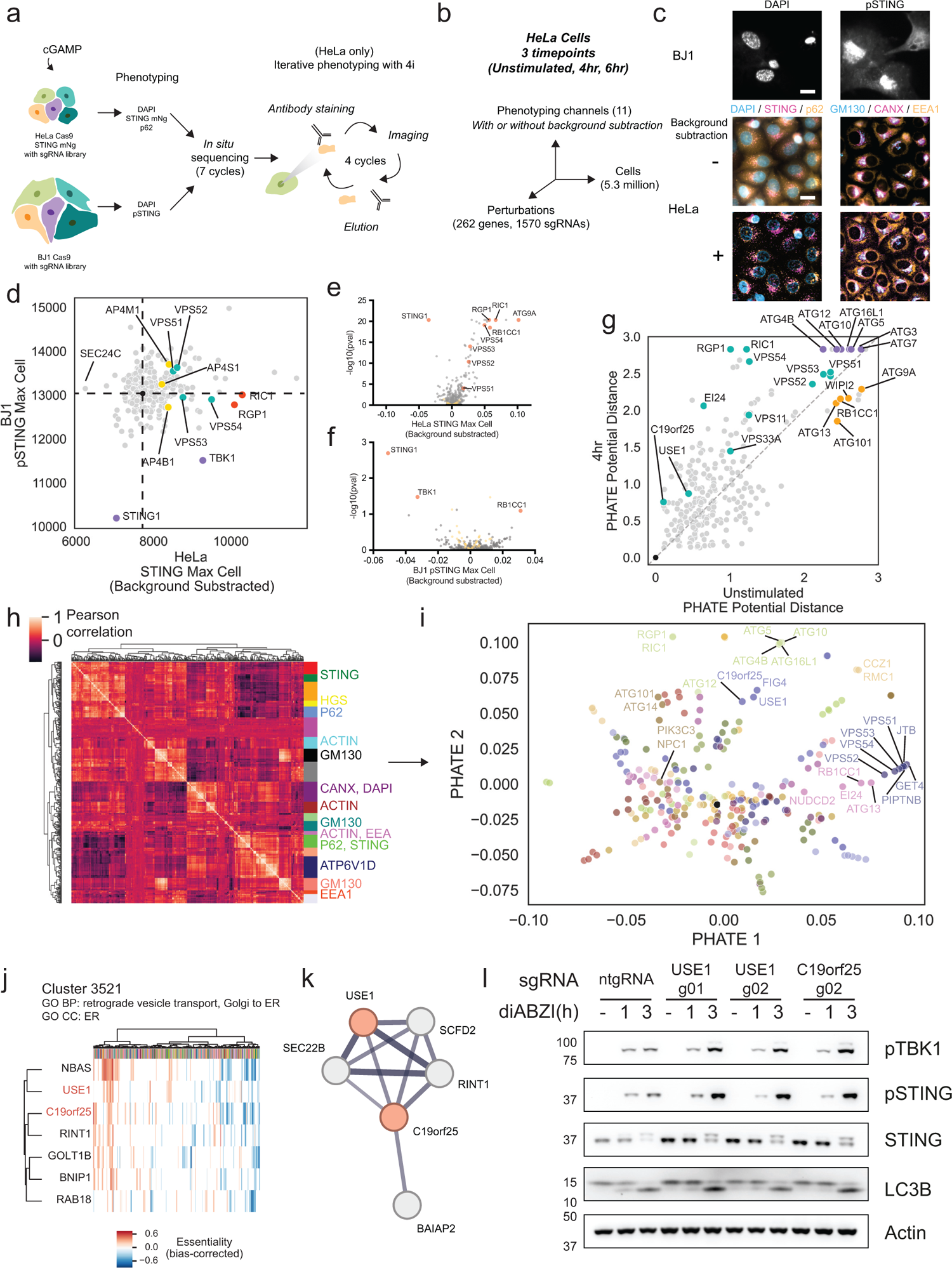
Secondary screens with additional dimensions further characterize genes regulating STING trafficking. **(A)** Workflow for targeted follow-up STING screens. **(B)** HeLa secondary screen is high-dimensional along multiple axes. **(C)** Selected fields of view from secondary screens for BJ1 cells and HeLa cells with or without background subtraction. Scale bar 20 µm. **(D)** Scatterplot of per-gene mean STING maximum (per cell) in HeLa cells compared to pSTING maximum per cell in BJ1 fibroblasts; black dot at the center of the dashed lines indicates non-targeting controls. Each point represents the average of measurements for each of the 262 targeted genes. **(E)** Volcano plot of mean STING maximum (per cell) in HeLa cells (deviation from ntgRNAs) and **(F)** pSTING maximum (per cell) in BJ1 fibroblasts (deviation from ntgRNAs); yellow dots indicate non-targeting control sgRNAs. Each point represents the average of measurements for the STING intensities on the x axis and the Stouffer aggregated p-values on the y axis each of the 262 targeted genes. **(G)** Correlation between phenotype (calculated as PHATE distance relative to NT sgRNAs) for each perturbed gene in the unstimulated vs 4 hour cGAMP-stimulated condition. **(H)** Hierarchical clustering based on Euclidean distance of HeLa screen features into 20 distinct modules. Clusters containing at least 40% of features related to a given channel are annotated. **(I)** PHATE dimensionality reduction and Leiden clustering of genes using as input mean per-gene scores across the 20 modules identified in (G). **(J)** Cluster 3521 and significant gene ontology (GO) terms as previously described^47^. X axis indicates distinct cell lines. C19orf25 and USE1 are manually highlighted. **(K)** STRING v11.5 physical subnetwork potential interactions (minimum score = 0.4) with C19orf25; USE1 and C19orf25 are manually highlighted. **(L)**. Immunoblot of the indicated proteins in control (ntgRNA) or USE1 or C19orf25 KO BJ1 fibroblasts stimulated with 1µM diABZI for the indicated times. One representative blot of n=3 experiments.

We identified factors affecting STING activation and degradation in both cell types based on pSTING and STING intensity (Fig. 3d**-f**). We then calculated the change in phenotype (relative to ntgRNAs) after dimensionality reduction of all features in cells that are unstimulated or after 4 hours of cGAMP treatment (Fig. 3g), aiming to prioritize genes with fewer morphological effects in the unstimulated condition. For example, we saw that factors involved in autophagy conjugation, a process required for both canonical autophagy and STING-induced CASM^44,45^, were more cGAMP-specific than upstream autophagy initiation factors required only for canonical autophagy (Fig. 3g). Next, we performed feature clustering to identify 20 modules of co-regulated features (Fig. 3h) in order to equalize the weights of feature types given the large number of channels and features in this screen, and performed PHATE dimensionality reduction and Leiden clustering on mean per-gene values across these feature modules (Fig. 3i**, S3c, Table S7**). This approach enables better distributed weighting of orthogonal feature groups.

By identifying genes in close proximity in the reduced dimensional space and focusing on perturbations that impact the stimulated more than the unstimulated phenotype (Fig. 3g), we observed that the poorly characterized gene C19orf25 co-clustered with USE1, a better-characterized ER membrane protein involved in Golgi-ER retrograde transport^46^ (Fig. 3i). USE1 was found proximal to STING in non-stimulated cells by proximity ligation proteomics^8^ (**Fig. S3d**). To confirm the association of C19orf25 with USE1 and STING, we examined clustering of co-essential modules from genome-wide fitness screens across cell types and observed again that C19orf25 was in the same cluster as USE1 in several instances, including cluster 3521, which contained genes related to retrograde Golgi-ER transport^47^ (Fig. 3j). Additionally, we used the STRING physical interaction subnetwork^48^ and observed that USE1 was one of five medium-confidence proteins that physically interacted with C19orf25 in immunoprecipitation assays (interaction score 0.582) (Fig. 3k). Analysis of deviation from ntgRNAs in the secondary screen showed that KO of both USE1 and C19orf25 led to increased STING colocalization with Golgi (GM130) and autophagy (p62) markers (**Fig. S3e**). Additionally, consistent with a role of USE1 in Golgi to ER recycling, KO of USE1 in non-stimulated cells led to an increase of STING colocalization with GM130, while KO of C19orf25 led to an increase in STING colocalization with Calnexin (CANX) (**Fig. S3f**), suggesting that these genes, similar to COPA^15–17,19^, play a role in STING Golgi-to-ER recycling. We further experimentally confirmed that knocking out C19orf25 and USE1 in BJ1 fibroblasts indeed leads to an increase in STING, pSTING, and pTBK1, consistent with the hypothesis that both genes are involved in retrograde Golgi-ER transport of proteins such as STING (Fig. 3l). Through optical pooled screening and integration of other datasets, we therefore inferred a biological function for C19orf25 and confirmed its role in regulation of STING activation.

We next focused on three complexes that have not been previously associated with STING trafficking and which have genetic mutations associated with disease in the human population: the HOPS complex, the GARP complex and the RIC1-RGP1 complex

### The HOPS complex subunits VPS11 and VPS33A regulate STING degradation at the lysosome

We next examined the impact of KO of the HOPS complex on STING trafficking and signaling, as the HOPS complex was not previously described to regulate STING degradation. This complex regulates membrane fusion in the endolysosomal compartment and has been shown to drive membrane fusion between autophagosomes and lysosomes^49^. Additionally, mutations in the VPS11 and VPS33A subunits of the complex have been found in patients with neurodegenerative diseases^50,51^.

Results from the genome-wide screen showed that the HOPS complex subunits VPS11, 33A, 18 and 16 clustered together (Fig. 2c) and increased STING max (max intensity of STING pixels) and mad (median absolute deviation of STING pixels) in the cytoplasm when knocked out (**Fig. S4a-b**). In the genome-wide screen, the ESCRT subunit VPS37A, which we showed regulates STING degradation in the endolysosomal compartment^8^, clustered with the HOPS complex (Fig. 2c**, 4a**), suggesting that the phenotype described in VPS37A KO cells could be similar to HOPS complex KO. The subunits in Cluster 9 of the genome-wide screen (Fig. 4a) are shared between the HOPS and the CORVET complex^52^. We excluded CORVET since VPS3 and VPS8 did not score in the genome-wide screen, while VPS39 and VPS41 were found to increase STING mad upon KO and VPS39 was found in Cluster 11 (Fig. 2c). To test if the HOPS complex plays a role in STING degradation, we knocked out VPS11 and VPS33A in a STING-mNeonGreen reporter line we previously described^8^ (**Fig. S4c**). This 293T reporter line expresses STING fused to mNeonGreen, allowing us to quantify STING degradation upon stimulation with the cGAMP stable homolog cGAMP(pS)2 via flow cytometry. KO of either subunit of the HOPS complex led to a reduction in STING degradation (Fig. 4b**-c**). We then tested the impact of VPS11 or VPS33A KO on STING signaling by knocking out either subunit with two independent sgRNAs in BJ1 fibroblasts. VPS11 or VPS33A KO led to a reduction in STING degradation, consistent with our screens and our reporter line, and increased STING phosphorylation both at 3 hours and 6 hours post-stimulation with cGAMP (Fig. 4d), resulting in increased expression of IFN-β and IL6 (Fig. 4e**, S4d**). As expected, KO of either HOPS subunit led to a defect in autophagic flux, as shown by accumulation of lipidated LC3B at steady state in absence of ligand (Fig. 4d). Interestingly, KO of either subunit also led to steady-state upregulation of the interferon-stimulated gene MX1, suggesting that defects in the HOPS complex drive spontaneous activation of type I IFN (Fig. 4d).

**Figure 4.**
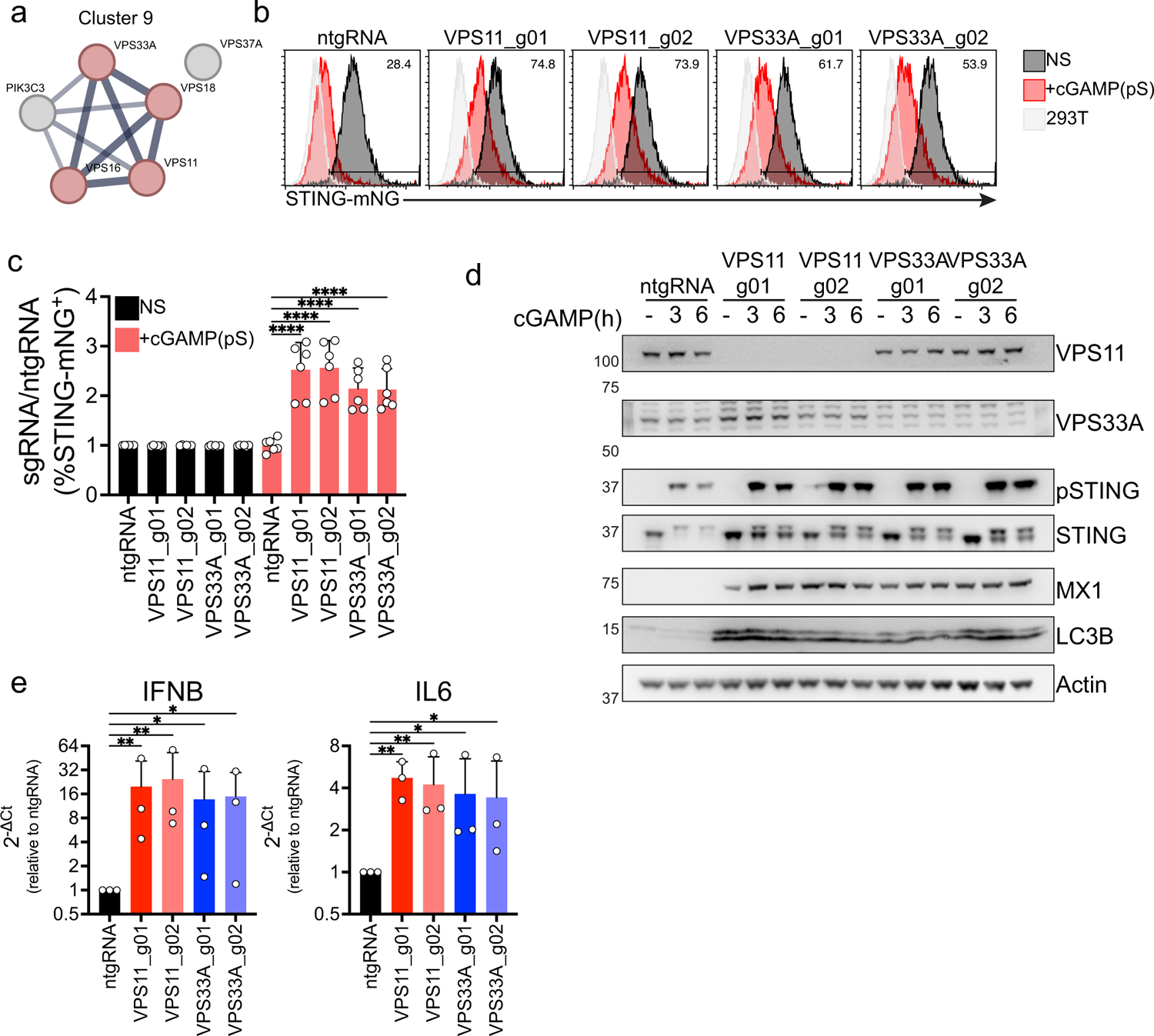
The HOPS complex subunits VPS11 and VPS33A are required for STING degradation. **(A)** STRING network of interaction of the indicated proteins found in Cluster 9 of the dimensionality reduced data shown in Figure 2. **(B)** mNG levels in 293 T STING-mNG cell lines KO for the indicated genes stimulated with 4μg/ml 2′3′-cGAMP(pS)2 (in medium) for 6h. One representative plot of n=3 independent experiments with n=2 technical replicates per experiment. **(C)** Percentage of STING-mNG positive cells in cells stimulated as in b. **(D)** Immunoblot of the indicated proteins in BJ1 fibroblasts transduced with a control guide (ntgRNA) or with VPS11 or VPS33A sgRNAs and stimulated with 0.5µg/ml cGAMP (in perm buffer) for the indicated times. **(E)** qPCR for IFNβ (left) and IL6 (right) in BJ1 fibroblasts KO for VPS11 (red) or VPS33A (blue) stimulated with 0.5 μg/ml cGAMP (in perm buffer) for 6h. n=3 independent experiments. 2^-ΔCt^ Fold Change calculated as ratio 2^-ΔCt^ sgRNA/2^-ΔCt^ ntgRNA for cells stimulated with cGAMP. One-way ANOVA on log-transformed data with Dunnet multiple comparison test. In all panels, bar plots show mean and error bars standard deviation. Marker unit for Western blots is KDa. *p <0.05,**p <0.01, ***p <0.001,****p < 0.0001, ns not significant.

Overall, we show that the HOPS complex subunits VPS11 and VPS33A regulate STING degradation. Blocking STING degradation results in increased phosphorylation of STING with consequent higher transcription of downstream IFN-β and IL6 and to upregulation of MX1 at steady-state.

### RIC1 and VPS52 regulate STING trafficking at the Golgi

We then characterized genes from two other complexes that increased per-cell STING maximum intensity and were identified by SVM classifiers and unsupervised clustering to affect STING trafficking (Fig. 2a**, S4b**). Specifically, we focused on the RAB6 GEF complex composed of RIC1 and RGP1 and the RAB6-interacting GARP complex^53,54^. Mutations in either RIC1 or subunits of the GARP complex have been associated with neurodegenerative disease in the human population. The knockout mouse for the GARP subunit VPS54, known as the Wobbler mouse, is a model of ALS^55–57^. In the genome-wide screen, RIC1 and RGP1 clustered together in cluster 0 and subunits of the GARP complex (VPS51, VPS52, VPS53, VPS54) clustered in cluster 67 with GET4, a protein involved in Golgi to ER trafficking and protein quality control^58^ and PITPNB, which is required for COPI-mediated retrograde transport from the Golgi^59^ (Fig. 5a). RIC1 was found enriched at 0.5 and 1h in the proximity of STING^8^ (**Fig. S5a**) and is part of the RIC1-RGP1 complex that regulates GDP-GTP exchange on RAB6, a RAB protein involved in the secretory pathway^60,61^. The VPS52 subunit of the GARP complex, a complex involved in endosome to Golgi recycling and membrane fusion at the Golgi, interacts with GTP bound RAB6^62^. Additionally, the constitutively active STING V155M found in SAVI patients has been shown to colocalize with RAB6 at steady-state^63^. We therefore hypothesized that both RIC1 and VPS52 regulate STING trafficking between the Golgi and the endosome.

**Figure 5.**
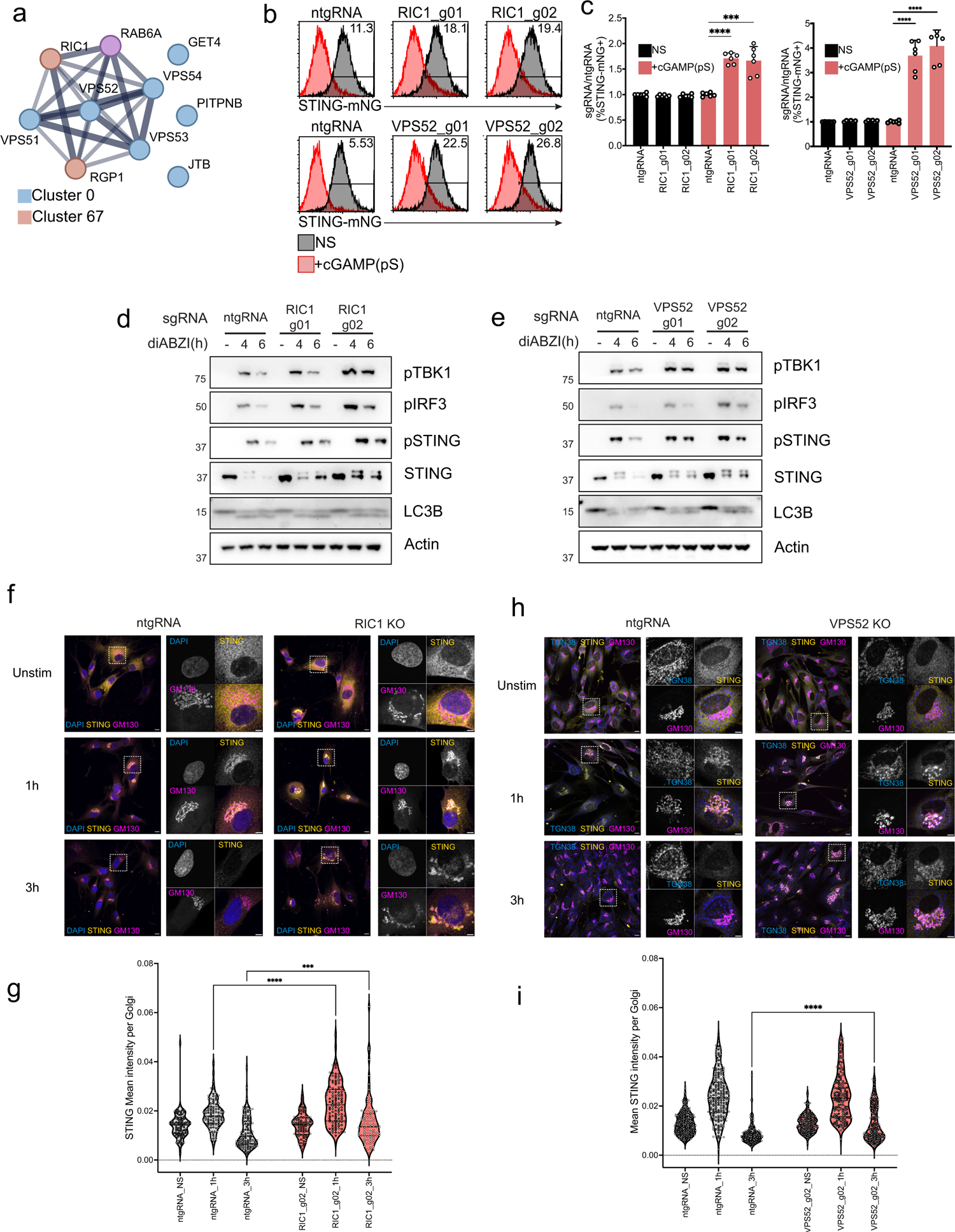
The RAB6 GEF RIC1 and the GARP complex subunit VPS52 regulate STING Golgi exit. **(A)** STRING network of interaction of the indicated proteins found in Cluster 9 of the dimensionality reduced data shown in Figure 2. **(B)** mNG levels in 293 T STING-mNG cell lines KO for the indicated genes stimulated with 4μg/ml 2′3′-cGAMP(pS)2 (in medium) for 6h. One representative plot of n=3 independent experiments with n=2 technical replicates per experiment. **(C)** Percentage of STING-mNG positive cells in cells stimulated as in B/ **(D)** Immunoblot of the indicated proteins in BJ1 fibroblasts transduced with a control guide (ntgRNA) or with RIC1 sgRNAs and stimulated with 1µM diABZI for the indicated times. **(E)** Same as in D for VPS52 KO cells. **(F)** Immunofluorescence of DAPI (blue), GM130 (magenta) and STING (yellow) in control (ntgRNA) or RIC1 KO BJ1 fibroblasts stimulated with 1µM diABZI for the indicated times. One field representative of n≥3 in n=3 independent experiments. **(G)** STING mean intensity calculated in GM130 (Golgi) regions in cells as in f. Each dot represents an individual Golgi. **(H)** Same as in e for VPS52 KO cells. **(I)** STING mean intensity calculated in GM130 (Golgi) regions in cells as in H. Each dot represents an individual Golgi. In all panels, bar plots show mean and error bars standard deviation. Marker unit for Western blots is KDa. *p <0.05,**p <0.01, ***p <0.001,****p < 0.0001, ns not significant.

We knocked out RIC1 or VPS52 in our STING-mNG reporter line (**Fig. S5b-d**). KO of either protein with two independent guides per gene led to a reduction in STING degradation (Fig. 5b-c), suggesting that both complexes play a role in STING trafficking. KO of either gene in BJ1 fibroblasts also led to a reduction in STING degradation upon stimulation, consistent with the reporter line, and led to an increase in pSTING, pTBK1 and pIRF3 downstream signaling while not affecting LC3B lipidation (Fig. 5d**-e**). These results suggest that RIC1 and VPS52 regulate STING signaling shutdown in addition to STING degradation, while they do not affect STING-dependent activation of autophagy or autophagy resolution. In order to validate the impact of each gene KO on STING trafficking at the Golgi, as predicted from our screens, we performed immunofluorescence in BJ1 fibroblasts that were either KO for RIC1 or VPS52. KO of each gene led to STING accumulation in the Golgi, shown by increased co-localization of STING and GM130 upon stimulation (Fig. 5f**-i**). Increased co-localization of STING and GM130 was particularly evident in RIC1 KO cells at 1 hour and either RIC1 or VPS52 KO led to an increase in colocalization at the 3 hour time-point, suggesting that both genes regulate STING exit from the Golgi. Increased Golgi dwell time in KO cells is consistent with an increase in pSTING levels due to a delay in STING trafficking and degradation (Fig. 5d**-e**). Overall, these data show that the RIC1-RGP1 complex and the VPS52 subunit of the GARP complex regulate STING trafficking and Golgi exit.

## Discussion

STING trafficking is central to innate responses driven by dsDNA of exogenous or endogenous origin. While activation of this pathway is essential to achieve effective antitumor responses, dysregulation of this process has been linked to disease. In order to identify genes regulating STING trafficking in an unbiased manner and facilitate the association of a genetic perturbation to a subcellular trafficking phenotype, we leveraged Optical Pooled Screens. We carried out one of the largest reported genome-wide OPS (over 45 million single-cell images assigned to genetic perturbations) and imaged cellular markers in 5 channels. We leveraged SVM classifiers to identify genes with statistically significant effects on STING trafficking and guide data interpretation, showing that this high-dimensional dataset performed better than previously performed CRISPR screens with flow cytometric or fitness readouts. Clustering of our high-content imaging dataset guided gene functional assignment. For example, we show that the recently characterized autophagy gene EI24^40^ plays a role in STING degradation and signaling shutdown based on its clustering neighbor ATG9A, which was previously known to regulate STING trafficking^41^.

To further confirm gene function assignment to phenotype, we performed targeted secondary OPS. To increase the information content of the screen, we integrated the iterative 4i staining protocol in the OPS pipeline, allowing us to image a total of 11 cellular markers. Additionally, we also performed for the first time an OPS in hTERT-immortalized primary cells, specifically BJ1 fibroblasts. Based on these datasets, we performed further clustering, which allowed us to assign a function to the poorly characterized gene C19orf25. C19orf25 clustered with USE1, a known regulator of Golgi-to-ER retrograde transport and we show that KO of C19orf25 phenotypically mimics USE1 KO on STING trafficking potentially through blocking STING Golgi-to-ER recycling.

By focusing on protein complexes that have mutations in the human population associated with disease but have not been shown to regulate STING trafficking, we also identified new roles for the HOPS, GARP and RIC1-RGP1 complexes in STING biology. We show that KO of the HOPS complex blocks STING degradation, similar to VPS37A KO that clustered with HOPS complex perturbations and that we previously demonstrated to regulate STING trafficking^8^. Similar to ESCRT KO or dysfunction, VPS11 and VPS33A KO lead to steady-state upregulation of the ISG MX1, suggesting that defects in the HOPS complex activate spontaneous type I IFN responses. STING vesicles are LC3B-lipidated post-Golgi via Conjugation of ATG8 on Single Membranes (CASM)^44,45^. The HOPS complex has an established role in driving fusion of autophagosomes with RAB7+ endolysosomes^49^. We speculate that the HOPS complex could help in fusion of STING-containing LC3B-lipidated vesicles with the endolysosome to achieve STING signaling shutdown. Under this hypothesis, defects in the complex would result in increased STING signaling. Interestingly, mutations in both VPS11 and VPS33A have been found in patients with neurodegenerative disease^50,51^. Exacerbated activation of STING via dysregulation of its trafficking has been associated with neurodegeneration in multiple diseases such as Niemann-Pick Disease and ALS^20,21,64^. The impact of HOPS complex KO on STING signaling opens to the possibility that dysfunction of this complex leads to exacerbated STING activation and subsequent pathogenesis in patients.

We also show that the RAB6 GEF complex composed of RIC1-RGP1 and the GARP complex subunit VPS52, which interacts with GTP-loaded RAB6, regulates STING exit from the Golgi. KO of either gene leads to a delay in Golgi exit of STING upon stimulation resulting in increased phosphorylation of STING and downstream TBK1 and IRF3. The GARP complex regulates endosome-to-Golgi recycling by promoting fusion of vesicles originating from the recycling endosome with the Golgi^62^. STING has been shown to traffic to the recycling endosome and STING-containing vesicles originating from this compartment are then degraded in an ESCRT-dependent manner^8–10^. Interestingly, in our proximity ligation proteomics, GO analysis showed enrichment of “Retrograde transport at the Trans-Golgi-Network” at 2 hours post STING stimulation^8^. Involvement of the GARP complex in STING trafficking opens the possibility that an endosome-to-Golgi recycling pathway could be present to recycle activated STING, analogous to Golgi to ER STING retrograde trafficking^15–17,19^.

Overall, we present here a collection of pooled screening results including a powerful high-content genome-wide OPS that constitutes a broad data resource for the immunology community. Integration with other results and further analysis may aid the identification of additional genes and mechanisms regulating STING trafficking and function. This data resource will also be of value to identify points of intervention in STING’s involvement in disease resulting from dysregulation of its trafficking or to boost STING responses in the context of antitumor immunity.

## Methods

### Cell lines

HeLa cells were described previously^23^ and had been transduced with a doxycycline-inducible Cas9 construct^28^. 293T were obtained from ATCC. 293T STING-mNeonGreen were described in Gentili et al 2023^8^. BJ-5ta (hTERT immortalized primary BJ fibroblasts - referred to as BJ1 in the text) were obtained from ATCC. HeLa and 293T were cultured in DMEM (Thermo Fisher) supplemented with 10% FBS (VWR), 1X Glutamax (Thermo Fisher) and 1% Penicillin/Streptomycin (Thermo Fisher). BJ-5ta were cultured in a 4:1 mix of DMEM (Thermo Fisher) and Medium 199 (Thermo Fisher) supplemented with 10% FBS (VWR), 1X Glutamax (Thermo Fisher) and 1% Penicillin/Streptomycin (Thermo Fisher). Cells were used up to passage 20 and were regularly tested for mycoplasma.

### Constructs

Doxycycline-inducible Cas9 was described previously^28^. For optical pooled screens, sgRNAs were cloned into the CROPseq-puro-v2 backbone (Addgene #127458). pXPR_BRD023 (lentiCRISPR v2) was a kind gift of the Broad GPP platform. sgRNAs for follow-up KOs were ordered as ssDNA oligos from Azenta and cloned in BsmBI digested pXPR_BRD023 with T4 DNA ligase (NEB). pTRIP-UbC-Blast-2A-STING-mNeonGreen was expressed in 293T STING-mNG and was described in Gentili et al 2023^8^. pTRIP-hPGK-Blast-2A-STING-mNeonGreen for expression in HeLa was obtained by substituting the UbC promoter for hPGK in pTRIP-UbC-Blast-2A-STING-mNeonGreen via PCR and Gibson assembly. psPAX2 (#12260) and pCMV-VSVG (#8454) were obtained from Addgene.

### Viral transductions for cell line generation

0.8 million 293T were seeded on the day of transduction in one well of a tissue culture treated 6 well plate. Cells were transfected with a mix containing 200µl OptiMEM (Thermo Fisher), 8µl TransIT-293 (Mirus Bio), 1µg psPAX2, 0.4µg pCMV-VSV-G and 1.6µg of construct DNA. 8µl TransIT were incubated in 200µl OptiMEM for 1 hour at room temperature (RT). DNA was then added to the mix, and the mix was shaken and incubated at RT for 30 minutes. 208µl/well of mix was then added to 293T. Medium was changed to 3ml of fresh 293T medium after overnight incubation. Virus supernatant was harvested 30-34 hours after medium change and filtered on 0.45µM syringe filters. 1 million 293T STING-mNG or 0.5 million BJ1 seeded the same day in one well of a 6 well plate were transduced with 2ml of virus including 8µg/ml Protamine (Thermo Fisher). Cells were incubated with viral supernatant for 1-2 days and cells were then transferred in a fresh flask for selection. Cells were then selected with the appropriate antibiotic: 2µg/ml Puromycin (Invivogen), 15µg/ml Blasticidin (Invivogen), 320µg/ml Hygromycin (Invivogen).

### Flow cytometry of 293T STING-mNG

For flow cytometry analysis of 293T STING-mNG, 0.15 million cells were seeded in tissue culture treated 24w plates the day before stimulation. Cells were then stimulated with 4µg/ml of cGAMP(pS)2 (Invivogen) for 6 hours. Cells were then lifted with TrypLE (Thermo Fisher), washed in medium and resuspended in MACS buffer (0.5% BSA, 2mM EDTA). Acquisition was performed on a Cytoflex S or Cytoflex LX (Beckman Coulter). Data were analyzed with FlowJo (BD).

### RT-qPCR

BJ1 fibroblasts were stimulated as described in figure legends. Cells were harvested and pellets were frozen at −80°C until processing. RNA was extracted using the Quick-RNA Microprep kit (Zymo) following manufacturer’s instructions. 5µl of RNA (roughly corresponding to 1µg) was reverse transcribed to cDNA using the LunaScript RT SuperMix kit - dye based qPCR detection (NEB) following manufacturer’s instructions. cDNA was diluted 1:2 with RNAse free water. 1µl of the reverse transcribed cDNA was used in each reaction for qPCR using the Luna Universal qPCR Master Mix (NEB) in a 10µl final reaction in a 384 well plate. 3 technical replicates per sample were measured on a CFX384 Touch Real-Time PCR Detection System (Biorad). Cycles were as follows: 1. Initial denaturation - 95°C, 60s; 2. Denaturation - 95°C, 15s; 3. Extension - 60°C, 30s; Melt Curve - 60-95°C. Steps 2-3 were repeated for 40 cycles. Primers were: IFNB1 FWD - CAGCATCTGCTGGTTGAAGA, RV - CATTACCTGAAGGCCAAGGA; IL6 FWD - CCCCTGACCCAACCACAAAT, RV - ATTTGCCGAAGAGCCCTCAG; GAPDH FWD - GTCTCCTCTGACTTCAACAGCG, RV - ACCACCCTGTTGCTGTAGCCAA. Data was analyzed using the 2^-ΔCt^ method relative to GAPDH.

### Western blot

Cell pellets were lysed in RIPA buffer (Boston Bioproducts) containing cOmplete, Mini, EDTA-free Protease Inhibitor Cocktail (Millipore Sigma) and PhosSTOP (Millipore Sigma) for 10 minutes on ice. Lysates were cleared by centrifugation at 16000g for 10 minutes at 4°C and Laemmli 6X, Sample buffer, SDS, Reducing (Boston Bioproducts) was added prior to loading. Samples were run on NuPAGE 4 to 12%, Bis-Tris Gels (Thermo Fisher) and transferred on nitrocellulose membrane with an iBlot2 (Thermo Fisher). Membranes were blocked in 5% non-fat milk in TBS Tween. Antibodies against phospho proteins were incubated in 5% BSA TBS tween. ECL signal was recorded on a ChemiDoc Biorad Imager. Data were analyzed with ImageLab (Biorad). Antibodies and dilutions are listed in Table S9.

### Immunofluorescence for follow-up studies

BJ1 were seeded on Fibronectin bovine plasma coated coverslips at 0.05 million cells/well density in a 24 well plate and stimulated as indicated in figure legends. Cells were then fixed with 2% Paraformaldehyde (Electron Microscopy Sciences) in PHEM buffer (Electron Microscopy Sciences) for 30 minutes at 37°C, washed three times with PBS and quenched with freshly prepared 0.1M Glycine for 10 minutes. Coverslips were then permeabilized and blocked with 10% goat serum (Thermo Fisher) in IF Perm Buffer (PBS, 0.5% BSA (Seracare), 0.05% Saponin from quillaja barka (Sigma)) for 20 minutes. Coverslips were then stained with primary antibodies for 1-2 hours at room temperature in IF Perm buffer supplemented with 10% goat serum, washed 5 times, and then stained with secondary antibodies in IF Perm Buffer for 1-2 hours. Coverslips were then washed 5 times, mounted with Fuoromont-G, with DAPI (Thermo Fisher) and dried at 37°C for one hour. Images were acquired on an Olympus IX83 using an Olympus PlanApo N 60X 1.42NA oil immersion objective controlled by Fluoview software.

### Image quantification for VPS52 and RIC1 Arrayed Knockout Experiments

CellProfiler version 4.2.5 was used to extract image-based features from 3 to 5 fields of view and three individual experiments per condition. GM130-positive areas were identified using the IdentifyPrimaryObjects module using the three-class Otsu thresholding method. Objects closer than 30 pixels were merged using the SplitOrMergeObjects module and the objects with an area smaller than 600 pixels were filtered out using the FilterObjects module. STING intensity within the GM130-positive objects was measured using the MeasureObjectIntensity module.

### OPS Library cloning, lentivirus production, transduction, and next generation sequencing of libraries

Libraries were cloned as previously described into a CROP-seq-puro-v2 (Addgene #127458) backbone. Lentivirus was then produced and transduced as previously described^23^. For library transductions, multiplicity of infection was estimated by counting colonies after sparse plating and antibiotic selection. Genomic DNA was also extracted for NGS validation of library representation. Genomic DNA was extracted using an extraction mix as previously described ^23^.. Barcodes and sgRNAs were amplified by PCR from a minimum of 100 genomic equivalents per library using JumpStart 2X Master Mix (initial denaturation for 5 minutes at 98°C, followed by 28 cycles of annealing for 10 s at 65°C, extension for 25 s at 72°C, and denaturation for 20 s at 98°C).

### STING stimulation, phenotyping, and in situ sequencing for genome-wide optical pooled screen

HeLa-TetR-Cas9 cells were transduced with pTRIP-hPGK-Blast-2A-STING-mNeonGreen and flow sorted twice to select cells with similar levels of reporter expression. Cells were maintained on blasticidin (20 μg/mL) for 4 days prior to transduction. For screening, cells were selected with puromycin (1 μg/mL) for 3 days after transduction and library representation was validated by NGS. Cas9 expression was induced with 1 μg/mL doxycycline for 1 week and cells were then seeded in six 6-well glass-bottom dishes at 415,000 cells/well 48 hours prior to fixation. cGAMP (Invivogen tlrl-nacga23) was added at 2 μg/mL for 10 minutes at 37°C in cGAMP Perm Buffer (50 mM HEPES (Corning), 100 mM KCl (Thermo Fisher), 3 mM MgCl2 (Thermo Fisher), 0.1 mM DTT (Thermo fisher), 85 mM Sucrose (Thermo Fisher), 0.5 mM ATP (Cayman chemicals), 0.1 mM GTP (Cayman Chemicals), 0.2% BSA (Seracare), 0.001% Digitonin (Promega)). 1mL warm media (DMEM with 10% FBS and 1% Pen-Strep) was then added to each well, media was quickly removed, and 2mL/well fresh media added for four hours prior to fixation. Cells were fixed by removing media and adding 4% paraformaldehyde (Electron Microscopy Sciences 15714) in PBS for 30 minutes at room temperature.

Cells were then permeabilized with 100% methanol for 20 minutes. The permeabilization solution was then carefully exchanged with PBS-T wash buffer (PBS + 0.05% Tween-20) by performing six 50% volume exchanges followed by three quick washes. The sgRNA sequence was reverse transcribed in situ overnight at 37°C using 1x RevertAid RT buffer, 250 μM dNTPs, 0.2 mg/mL BSA, 1 μM RT primer with LNA bases -A+CT+CG+GT+GC+CA+CT+TTTTCAA -, 0.8 U/μL Ribolock RNase inhibitor, and 4.8 U/μL RevertAid H minus reverse transcriptase in 750 μL/well as previously described^23^. After reverse transcription, cells were washed 5x with PBS-T and post-fixed using 3% paraformaldehyde and 0.1% glutaraldehyde in PBS for 30 minutes, followed by washing with PBS-T 3 times.

Cells were stained with primary antibodies for GM130 (1:1300, Cell Signaling Technology Cat# 12480, RRID:AB_2797933) and p62 (1:300, Cell Signaling Technology Cat# 88588, RRID:AB_2800125) in 3% BSA (VWR Cat# 97061-422) in PBS for 4 hours. Samples were then washed 3x in PBS-T for 3 minutes, and incubated with secondary antibodies: 1:1800 donkey anti-mouse antibody (Jackson ImmunoResearch Labs Cat# 715-006-151, RRID:AB_2340762) disulfide-linked to Alexa Fluor 594 (Thermo Fisher A10270) via a custom conjugation, 1:1800 donkey anti-rabbit antibody (Jackson ImmunoResearch Labs Cat# 711-006-152, RRID:AB_2340586) disulfide-linked to Alexa Fluor 647 (Thermo Fisher Scientific A10277) via a custom conjugation, in 3% BSA for 2 hours at 37°C. Finally, cells were washed 6x in PBS-T, and stained with BV421-conjugated anti-CD63 (1:1000, BioLegend Cat# 353029, RRID:AB_2687003). Cells were then imaged and Alexa Fluor 594 and 647 antibodies were destained with 50mM TCEP in 2x SSC for 45 minutes at room temperature. Subsequently, samples were incubated in a padlock probe and extension-ligation reaction mixture (1x Ampligase buffer, 0.4 U/μL RNase H, 0.2 mg/mL BSA, 100 nM padlock probe - /5Phos/GTTTCAGAGCTATGCTCTCCTGTTCGCCAAATTCTACCCACCACCCACTCTCCAAAGGACGAAACACCG -, 0.02 U/μL TaqIT polymerase, 0.5 U/μL Ampligase and 50 nM dNTPs) for 5 minutes at 37°C and 90 minutes at 45°C, and finally washed 2 times with PBS-T. Circularized padlocks were amplified using a rolling circle amplification mix (1x Phi29 buffer, 250 μM dNTPs, 0.2 mg/mL BSA, 5% glycerol, and 1 U/μL Phi29 DNA polymerase) at 30°C overnight. DAPI (600 ng/mL) was then added to cells and cells were imaged in the DAPI and STING mNeonGreen channels for alignment to the first round of phenotyping images. Finally, in situ sequencing was performed as previously described using the sequencing primer GCCAAATTCTACCCACCACCCACTCTCCAAAGGACGAAACACCG for 12 cycles.

### Secondary optical pooled screens

Genes to target for the secondary screen were manually chosen based on ranking in the optical pooled screen SVMs, the meta-analysis by information content rankings, and adding select genes of interest from the TurboID dataset^8^. hTert-BJ1 (BJ-5ta - CRL-4001) cells from ATCC were used for secondary screening along with HeLa-TetR-Cas9 cells described above.

Prior to screening, BJ1 cells were transduced with Cas9 (Addgene #96924) and selected using 10 µg/mL blasticidin for 7 days. Subsequently, library transduction was performed as described for HeLa cells above without the addition of doxycycline. BJ1 cells were plated for stimulation 5 days post-transduction at 150,000 cells/well and stimulated with cGAMP and buffer at 7 days post-transduction as described above for HeLa cells.

For BJ1 secondary screening, in situ sequencing and antibody staining were performed as described for the HeLa cells but only imaged for one round of immunofluorescence: 200 ng/mL DAPI with 1:100 pSTING Ser366 (Cell Signaling Technology Cat# 19781, RRID:AB_2737062) as the primary antibody detected with 1:900 custom-conjugated donkey anti-rabbit antibody (Jackson ImmunoResearch Labs Cat# 711-006-152, RRID:AB_2340586) disulfide-linked to Alexa Fluor 647 (Thermo Fisher Scientific A10277).

For HeLa cell secondary screening, one round of antibody staining was performed after reverse transcription and post-fixation, using p62 mouse (1:250, Cell Signaling Technology Cat# 88588, RRID:AB_2800125) for two hours followed by 1:900 anti-mouse antibody (Jackson ImmunoResearch Labs Cat# 715-006-151, RRID:AB_2340762) disulfide-linked to Alexa Fluor 594 (Thermo Fisher A10270) via a custom conjugation for 1 hour and imaged in 200ng/mL DAPI in 2X SSC. Cells were then imaged and the Alexa Fluor 594 antibody was destained with 50mM TCEP in 2x SSC for 45 minutes at room temperature. Subsequently, samples were incubated in a padlock probe and extension-ligation reaction mixture and amplified using rolling circle amplification as described above. Following 7 cycles of in situ sequencing, HeLa cells were iteratively assayed using 4i^42^ for 4 additional imaging cycles (see below):

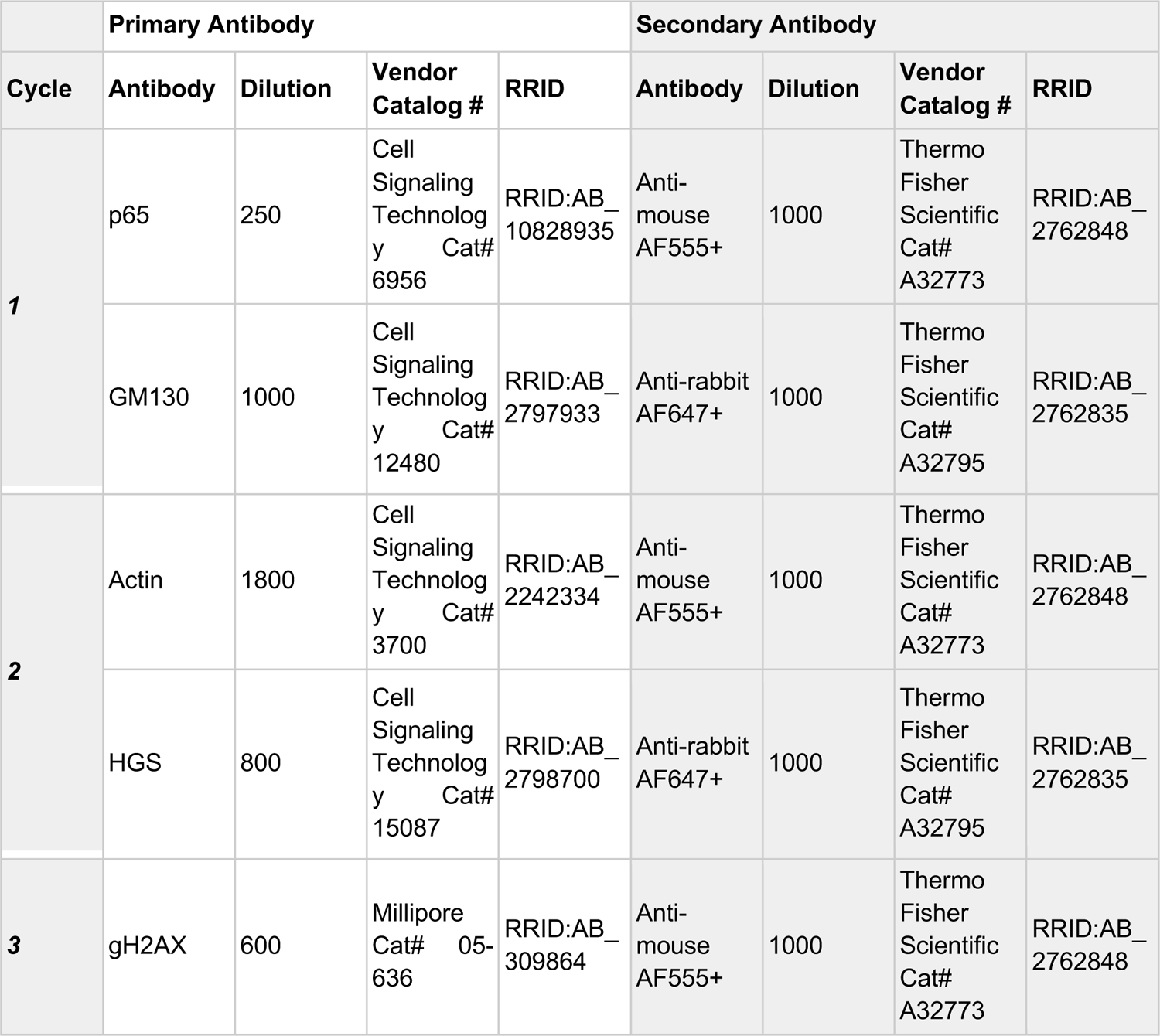
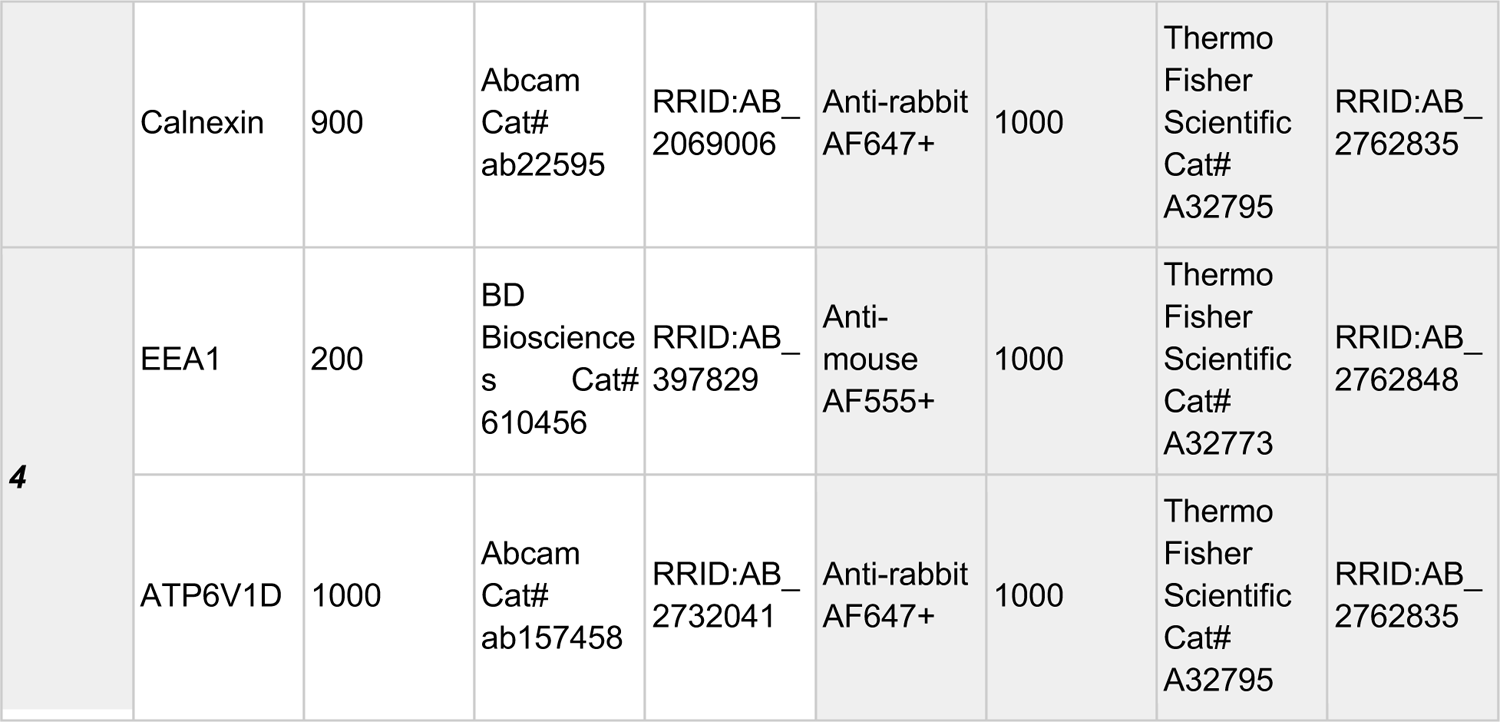

To perform iterative fluorescence, after antibody staining and imaging, stripping of fluorescence signal was performed similarly to what was previously described^42^. Briefly, after imaging samples were washed six times quickly in distilled water. Water was then carefully aspirated and elution buffer (previously described) was added to the sample with shaking at 300 rpm for 30 minutes. Samples were imaged in imaging buffer as previously described^42^ with the addition of 50 mM HEPES and pH was adjusted to 7.4^65^.

### Fluorescence microscopy

All in situ sequencing images were acquired using a Ti-2 Eclipse inverted epifluorescence microscope (Nikon) with automated XYZ stage control and hardware autofocus. The Lumencor CELESTA Light Engine was used for fluorescence illumination and all hardware was controlled using NIS elements software with the JOBS module. In situ sequencing cycles were imaged using a 10X 0.45 NA CFI Plan Apo λ objective (Nikon MRD00105) with the following filters (Semrock) and exposure times for each base: G (546 nm laser at 40% power, emission 575/30 nm, dichroic 552nm, 200 ms); T (546 nm laser at 40% power, emission 615/24 nm, dichroic 565 nm, 200 ms); A (637 nm laser at 40% power, emission 680/42 nm, dichroic 660 nm, 200 ms); C (637 nm laser at 40% power, emission 732/68 nm, dichroic 660 nm, 200 ms).

For the genome-wide primary screen, phenotyping images were acquired using a 20X 0.75 NA CFI Plan Apo λ objective (Nikon MRD00205) with the following filters (Semrock unless otherwise noted) and exposure times: CD63 (405 nm laser at 10% power, Chroma Multi LED set #89402, 30ms), DAPI (405 nm laser at 5% power, Chroma Multi LED set #89402, 10ms), STING mNeonGreen (477 nm laser at 30% power, Chroma Multi LED set #96372, 200ms), AF594 (546 nm laser at 80% power, emission 615/24 nm, dichroic 565 nm, 400ms), AF647 (637 nm laser at 30% power, emission 680/42 nm, dichroic 660 nm, 100ms). Two z-stacks 1.5 µm apart were acquired and images were max-projected.

For the secondary screen, phenotyping images for HeLa cells were acquired using a 40x 0.95 NA CFI Plan Apo λ objective (Nikon MRD70470) with the following filters and exposure times: DAPI (405 nm laser at 100% power, Chroma Multi LED set #89402, 100ms), AF594 p62 (546 nm laser at 100% power, Chroma Multi LED set #89402, 100ms), and STING mNeonGreen (477 nm laser at 100% power, Chroma Multi LED set #96372, 100ms). Subsequent rounds after in situ sequencing were imaged for DAPI, AF555, and AF647 with the same filter sets and laser lines noted above for DAPI, AF594, and AF647, respectively, with the following laser powers and exposure times at 100% power: Cycle 1 - 1/10/10ms; Cycle 2 - 1/5/10/ms; Cycle 4 - 1/30/30ms; Cycle 4 - 4/20/30ms.

Phenotyping images for BJ1 cells were acquired using a 20X 0.75 NA CFI Plan Apo λ objective (Nikon MRD00205) with the following filters and exposure times: DAPI (405 nm laser at 100% power, Chroma Multi LED set #89402, 4ms), and AF647 (637 nm laser at 100% power, Chroma Multi LED set #89402, 40ms). Two z-stacks 1.5 µm apart were acquired and max-projected.

### Phosphorylated STING flow cytometry screen

See Liu, Carlson et al. Science 2023^12^. RFP-LC3B FIP200 KO 293T cells were transduced with pTRIP-PGK-Blast-P2A-STING-HA and selected with 20 µg/mL blasticidin (Thermo Fisher Scientific #A1113903). For screening, 200M RFP-LC3+ STING-HA+ FIP200 KO 293T cells were transduced with Cas9-sgRNA all-in-one Brunello library at MOI=0.4 and selected with 2 µg/mL puromycin for 2 days. 8 days after transduction, 200M transduced cells were treated with 1 µM diABZI (Invivogen, #tlrl-diabzi) for 2.5hr, and then permeabilized with 1X perm buffer (PBS with 0.05% saponin and 0.1% glycine) for 4 min (18). Permeabilized cells were then washed with 1X PBS, and fixed, permeabilized further, and stained using BD Cytofix/Cytoperm kit (#554714) following staining with anti-Phospho-STING (Ser366) (D8K6H) Rabbit mAb (Alexa Fluor® 488 Conjugate) (CST, #41622) and anti-HA Alexa647 (Biolegend, #682404) at 1:200 dilution.

### Quantification and Statistical Analysis Image analysis

Images of cellular phenotype and in situ sequencing of perturbations were manually aligned during acquisition using nuclear masks to calibrate the plate position to each of the four corner wells of a 6-well plate. For the genome-wide screen, images between phenotyping cycles were aligned using masks generated using the STING mNeonGreen channel. Alignment was subsequently refined computationally via cross-correlation of DAPI signal between imaging acquisitions. Nuclei and cells were detected and segmented as previously described and in situ sequencing read calling was performed as previously described^66^. Data analysis functions were written in Python, using Snakemake for workflow control^67^. Image analysis code is available on GitHub. Briefly, for segmentation of phenotyping images from the primary screen, nuclei were segmented using the following parameters: using the following parameters: nuclei smooth = 1.15, nuclei radius = 15, nucleus area 110-1600, DAPI intensity threshold = 6100. Cells were segmented using signal in the GM130 channel, at an intensity threshold = 2800. For segmentation of in situ sequencing images from the primary and secondary screens, nuclei were segmented using the following parameters: nuclei smooth = 1.15, nuclei radius = 15, nucleus area 20-800 (BJ1), 20-400 (HeLa), DAPI intensity threshold = 1500 (BJ1), 2000 (HeLa). Cells were segmented using signal in the four sequencing channels at intensity thresholds adjusted for each plate, between 3800 and 4200. For segmentation of phenotyping images from the secondary screen, nuclei were segmented using the following parameters: nuclei smooth = 25 (BJ1), 9 (HeLa), nuclei radius = 30 (BJ1), 100 (HeLa), nucleus area 700-180000 (BJ1), 200-18000 (HeLa), DAPI intensity threshold = 1900 (BJ1), 1400 (HeLa). All other parameters used for analysis were set to default settings. Background subtraction using a rolling ball algorithm was performed with radius = 50 for all channels in the secondary screen.

### Optical pooled screen analysis

Only cells with a minimum of one read matching a barcode in the library were analyzed. For the genome-wide screen, only genes with a minimum of one read matching an sgRNA in the library and 2 sgRNAs with at least 50 cells/sgRNA were considered for analysis. Features were normalized on a per-cell basis relative to cells in the same field of view by subtracting the median and dividing by the MAD x 1.4826^68^. SVM models were trained using the Intel Extension for Scikit-learn^69^ and the command svm.LinearSVC(random_state=7, C=1e-5, max_iter = 100000, tol = .00005, class_weight = ‘balanced’) with an 80/20 train/test split for the perturbed classifier and using all cells for each sgRNA in the unstimulated and increased stimulation classifiers, as these two classifiers were separately pre-trained on non-targeting control cells with the same SVM parameters as the perturbed classifier. Resulting mean model scores were averaged over sgRNAs for a given gene and significance was determined by comparing scores for individual sgRNAs to distributions bootstrapped from non-targeting control cells (bootstrapped 100,000 times). Gene-level p-values were calculated using Stouffer’s method to aggregate across sgRNAs targeting the same gene and subsequently corrected using the Benjamini-Hochberg procedure.

### Meta-Analysis by Information Content

MAIC scores were calculated for each screen (Table S3) using MAIC version 1.0 with default parameters^26^). Pearson correlations of MAIC scores between different screens are shown in Fig 1i. For each screen, the top 500 genes in each direction (if applicable) significant at p < 0.05 were selected as input except for the TurboID dataset, where the full list of 132 genes significant at p < 0.07 was used as discussed previously^8^.

### Dimensionality reduction and clustering

PHATE 1.0.10^33^ was used to cluster genes using mean gene-level feature values. For the genome-wide screen, PCA was performed and 31 PCs were selected, preserving 95% of variance in the data. PHATE was then performed using the following parameters: knn_dist = ‘euclidean’, gamma = 1, knn = 2, random_state = 7). For the secondary screen, PHATE was performed using no PCA prior to clustering, since the dataset was already reduced to 20 features by co-clustering features using hierarchical clustering and taking average values across all features in each cluster. The following parameters were then used for PHATE clustering: knn_dist = ‘euclidean’, gamma = 0, knn = 1, random_state = 7, mds_dist = ‘cosine’. Leiden clustering was also performed. To determine the optimal number of clusters, Leiden clustering was performed at varying resolutions on 90% of the data (selected across 15 iterations) and cluster stability assessed using ARS (sklearn.metrics.adjusted_rand_score)^70^ in order to select an optimal resolution parameter for clustering. To identify clusters enriched for hits from our classifiers or significant DEGs from the genome-scale Perturb-seq data, we used the rank genes groups function in scanpy^71^.

### Phosphorylated STING flow cytometry analysis

Guide RNA abundances were extracted from FASTQ files using poolq 3.3.2 with fixed row and barcode policies. Resulting log-abundances were then subtracted between sorting bins from each experimental condition and analyzed using the Broad Genetic Perturbation Platform screen analysis tool (https://portals.broadinstitute.org/gpp/public/analysis-tools/crispr-gene-scoring) using a hypergeometric analysis.

## Data and Code Availability

Code is available at https://github.com/beccajcarlson/STINGOpticalPooledScreen. Imaging data is available on Google Cloud Storage at gs://opspublic-east1/STINGOpticalPooledScreen.

## Acknowledgements

We thank members of the Blainey and Hacohen labs for critical feedback and discussions. We thank Celeste Diaz and Julia Bauman in the lab of J.T. Neal at the Broad Institute for assistance in developing custom antibody conjugations. The HeLa cell line was used in this research. Henrietta Lacks, and the HeLa cell line that was established from her tumor cells without her knowledge or consent in 1951, have made significant contributions to scientific progress and advances in human health. We are grateful to Lacks, now deceased, and to the Lacks family for their contributions to biomedical research. This work was supported by the Broad Institute through startup funding (to P.C.B.) and the BN10 program, two grants from the National Human Genome Research Institute (HG009283 and RM HG006193) and one grant from NIH to N.H. (1R01AI158495). P.C.B. was supported by a Career Award at the Scientific Interface from the Burroughs Wellcome Fund. R.J.C. is supported by a Fannie and John Hertz Foundation Fellowship and an NSF Graduate Research Fellowship. M.G. was supported by an EMBO Long-Term Fellowship (ALTF 486-2018) and is a Cancer Research Institute/Bristol Myers Squibb Fellow (CRI 2993). Y.Q. is supported by the NIH under grant K00 CA264422 and by funding from the Eric and Wendy Schmidt Center at the Broad Institute of MIT and Harvard.

## Author Contributions

M.G., R.J.C., and B.L. designed the approach with input from all authors. M.G., R.J.C., B.L., J.A., and Q.H. performed experiments. R.J.C., Y.Q. M.G., and Q.H. analyzed data. P.C.B. and N.H. supervised the research. M.G. and R.J.C. wrote the manuscript with contributions from all authors.

## Supplementary Information

**Table S1.** Genome-wide optical pooled screen mean per-gene features.

**Table S2.** Genome-wide OPS SVM classifier results.

**Table S3.** Meta-analysis by information content results.

**Table S4.** Dimensionality reduction clusters from genome-wide screen.

**Table S5.** Secondary screen mean per-gene features for HeLa cells.

**Table S6.** Secondary screen mean per-gene features for BJ1 cells.

**Table S7.** Secondary screen dimensionality reduction results and unstimulated vs 4 hour PHATE potential distances.

**Table S8.** List of sgRNAs used in the study

**Table S9.** List of antibodies used in the study

## Supplementary Figures 1-5

**Figure S1.**
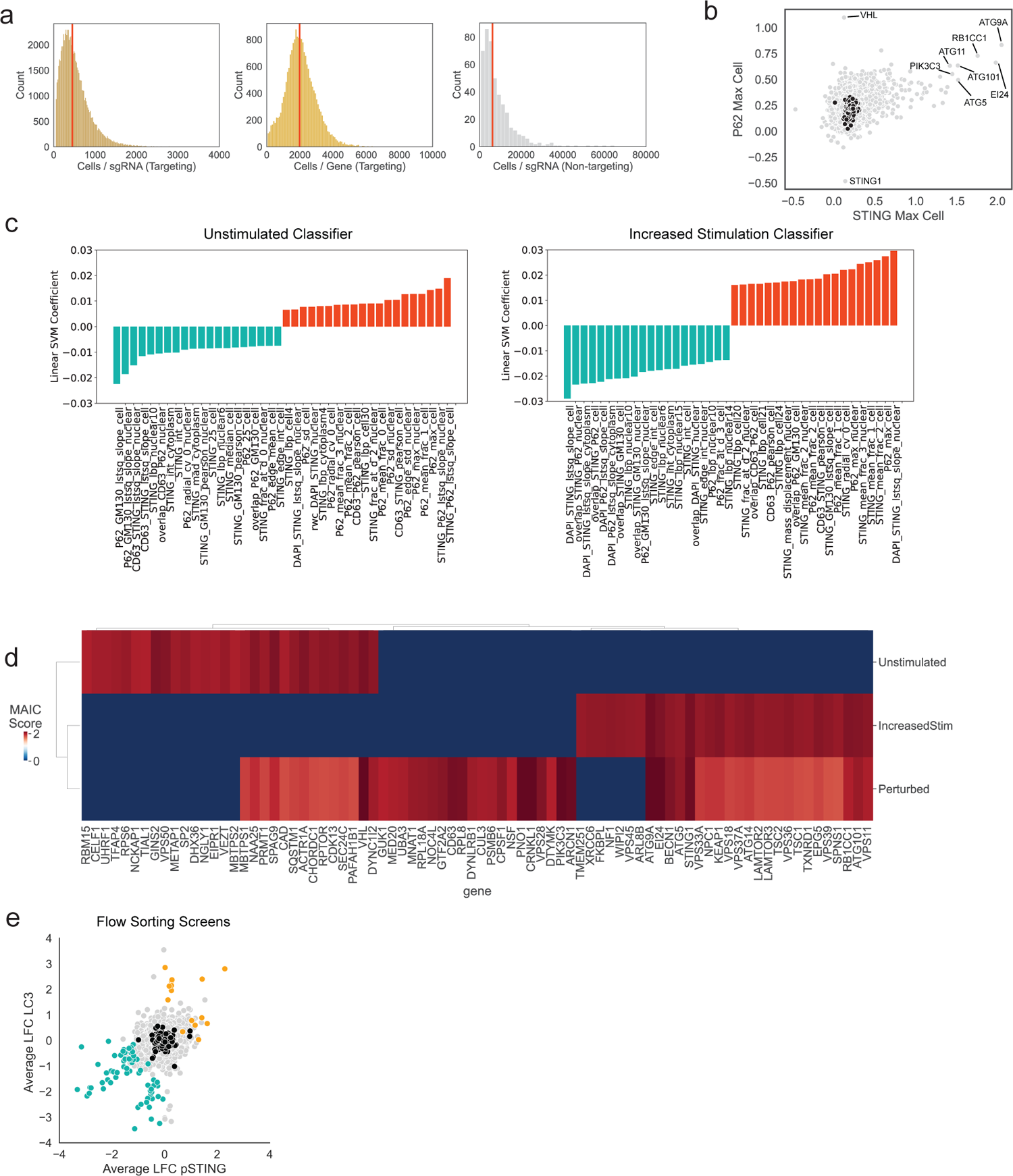
**(A)** Distribution of number of cells in the screen for each sgRNA or gene for targeting sgRNAs and non-targeting controls. Medians highlighted in red. **(B)** Scatterplot of STING per-cell maximum intensity and p62 maximum intensity for each gene in the screen. Black dots indicate non-targeting control sgRNAs. **(C)** Top and bottom 20 feature weights for SVM unstimulated and increased stimulation classifiers. **(D)** MAIC scores for top 30 genes (by overall MAIC score) for each OPS SVM classifier. **(E)** Correlation of log2 fold change (LFC) for pSTING and LC3 in STING flow cytometry screens. Orange: genes that increased both LC3 and pSTING at p < .001 in both screens; blue: genes that decreased both metrics at the same significance.

**Figure S2.**
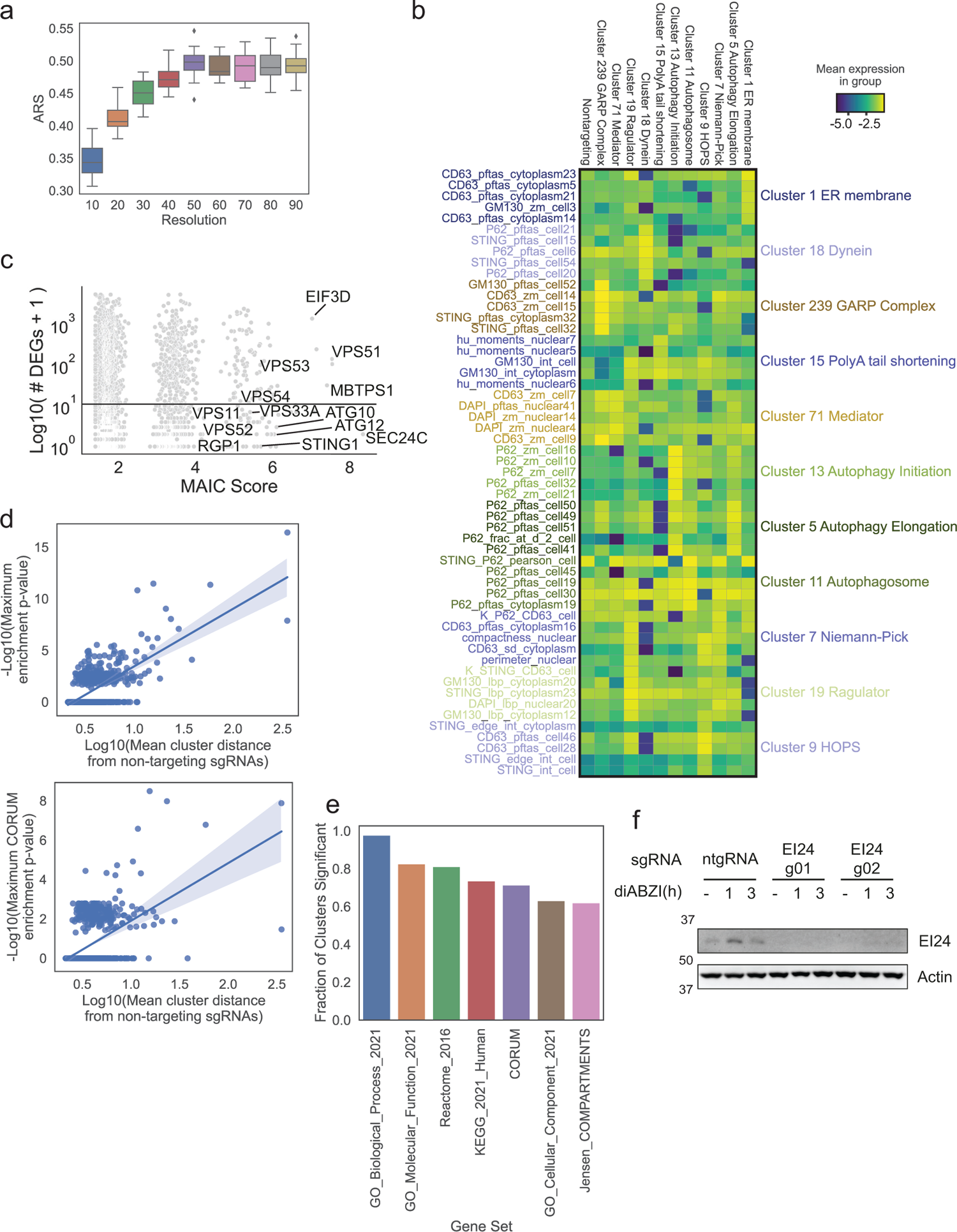
**(A)** Adjusted Rand score for Leiden clustering at different resolutions. **(B)** Top 5 features significantly differentiating clusters highlighted in Figure 2. **(C)** MAIC Score plotted against number of DEGs for genes included in the genome-scale Perturb-seq K562 dataset. **(D)** Scatterplots of mean cluster distance from non-targeting sgRNAs (PHATE potential distance) against the maximum enrichment p-value from Enrichr (GO, Reactome, KEGG, Jensen COMPARTMENTS and CORUM datasets, top) or from CORUM alone (bottom) **(E)** Fraction of clusters (among clusters with >1 gene and >20% of genes not non-targeting sgRNAs) that had at least one significant term as calculated by Enrichr for the noted categories. **(F)** Immunoblot of the indicated proteins in BJ1 fibroblasts transduced with a control guide (ntgRNA) or with EI24 sgRNAs and stimulated with 1µM diABZI for the indicated times.

**Figure S3.**
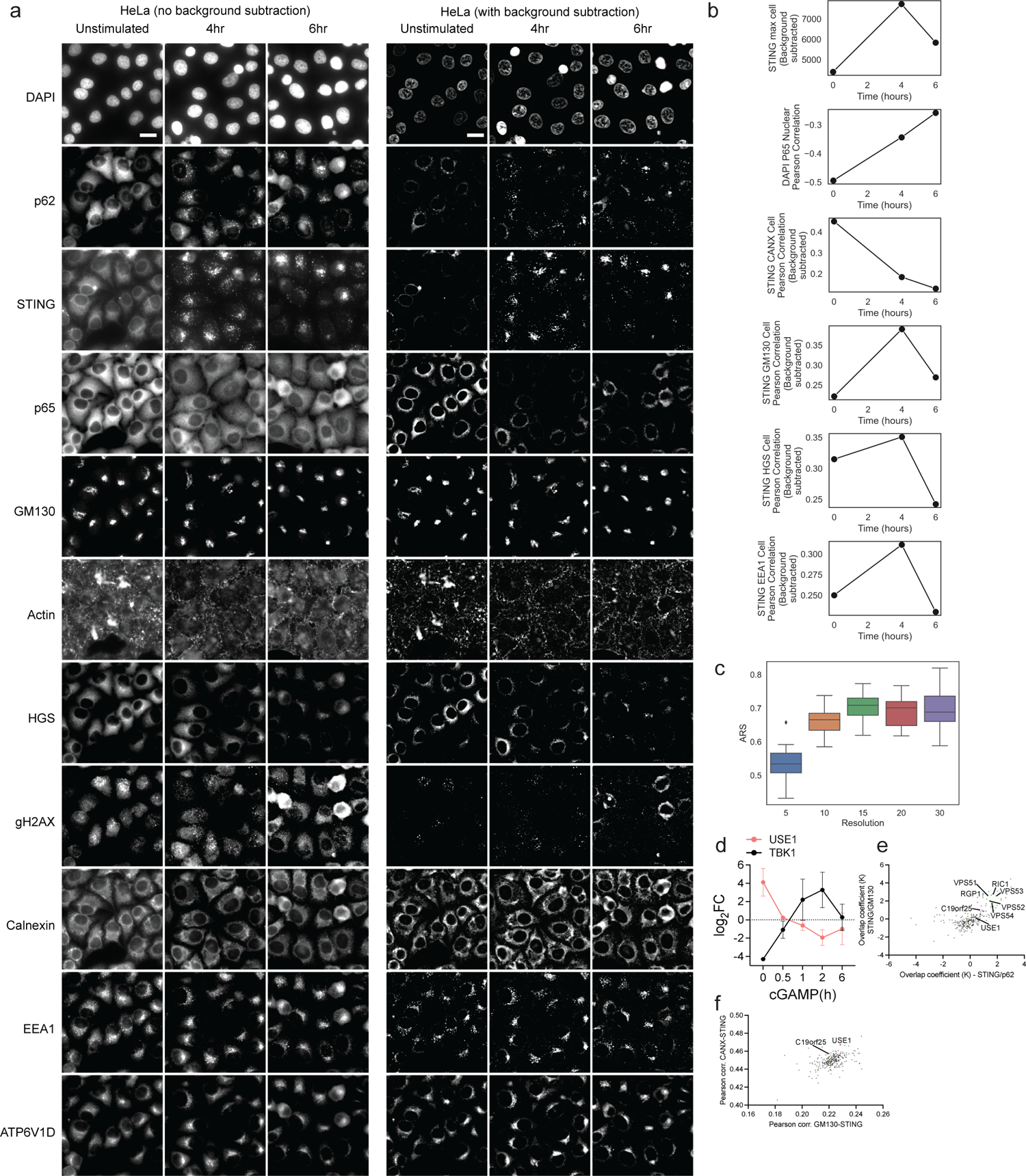
**(A)** Selected fields of view from secondary screens for HeLa cells, all channels shown. Scale bar 20 µm. **(B)** HeLa secondary screen non-targeting mean feature values across time. **(C)** Adjusted Rand score (ARS) for Leiden clustering at different resolutions. **(D)** log2 fold change (log2FC) enrichment of the indicated proteins in the STING-TurboID datasets at the indicated timepoints post cGAMP stimulation. **(E)** Overlap coefficient for the indicated channels in integrated deviation from ntgRNA from secondary OPS. **(F)** Pearson correlation of the indicated channels in unstimulated cells from the secondary OPS.

**Figure S4.**
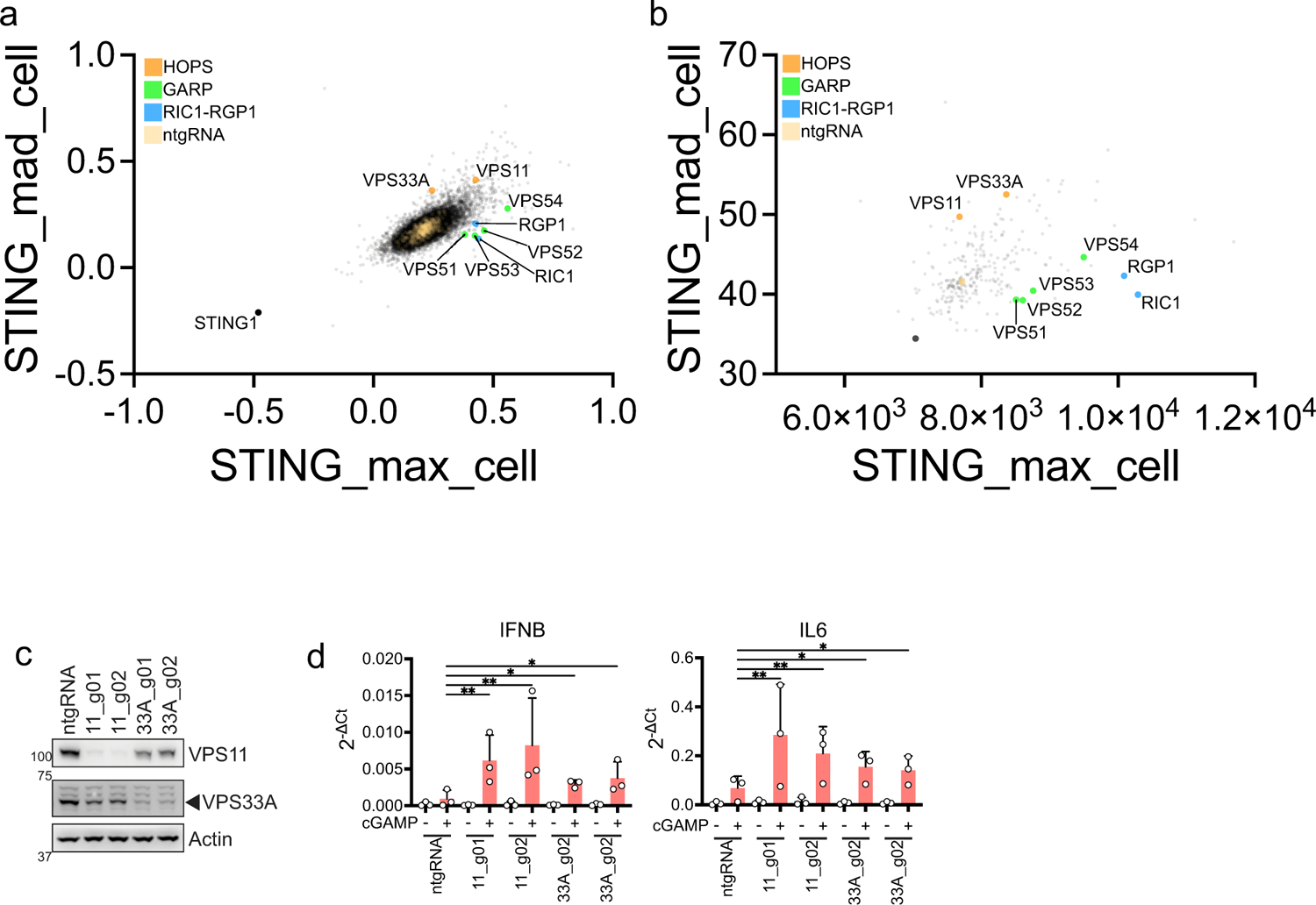
**(A)** STING_max and STING_mad extracted features correlation in the genome-wide OPS in HeLa cells. Specific subunits of complexes of interest are indicated in color. STING1 is highlighted in black. Non-targeting control sgRNAs are indicated in yellow. Pixel intensity is normalized to ntgRNAs. **(B)** STING_max and STING_mad extracted features correlation in the genome-wide OPS in HeLa cells. Specific subunits of complexes of interest are indicated in color. STING1 is highlighted in black. Non-targeting control sgRNAs are indicated in yellow. Background subtracted pixel intensity values are plotted. **(C)** Immunoblot of the indicated proteins in 293T STING-mNG transduced with control (ntgRNA) or VPS11 or VPS33A sgRNAs. One blot representative of n=3 blots. **(D)** Raw 2^-ΔCt^ values related to Fig. 4e.

**Figure S5.**
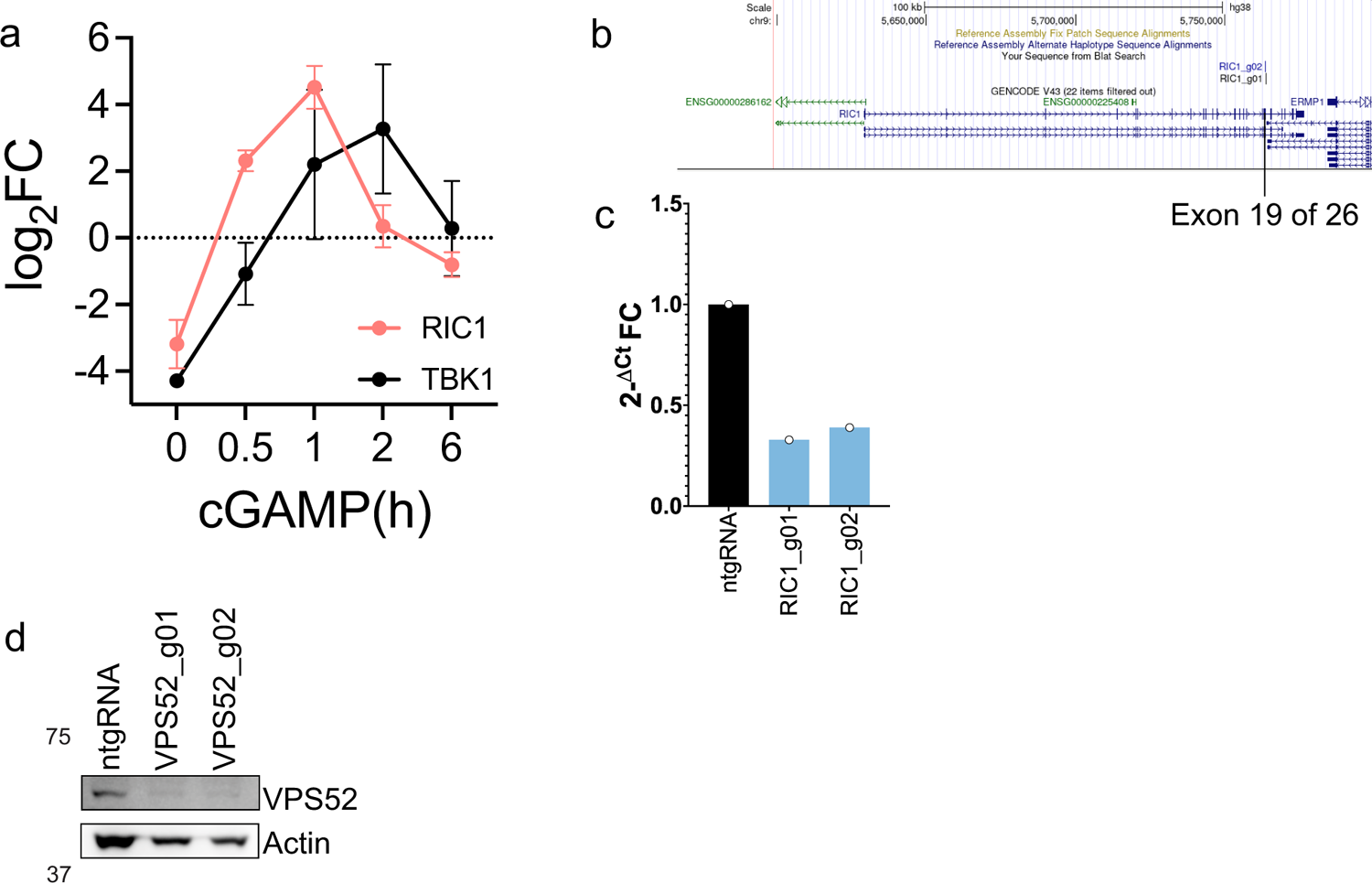
**(A)** log2 fold change (log2FC) enrichment of the indicated proteins in the STING-TurboID datasets at the indicated timepoints post cGAMP stimulation. **(B)** Location of the two RIC1 targeting sgRNAs used in this paper visualized in UCSC Genome browser. While we could not identify a reliable antibody for RIC1, both guides targeted the gene in exon 19 of 26 at >50-55nt from the exon-exon junction potentially triggering Non-Sense Mediated Decay^72^. **(C)** qPCR of RIC1 expression in 293T STING-mNG transduced with a control (ntgRNA) or RIC1 targeting guides as in b). **(D)** Immunoblot of the indicated proteins in 293T STING-mNG transduced with a control (ntgRNA) or VPS52 targeting sgRNAs.

